# Reciprocal regulation of pancreatic ductal adenocarcinoma growth and molecular subtype by HNF4α and SIX1/4

**DOI:** 10.1101/814525

**Authors:** Soledad A. Camolotto, Veronika K. Belova, Luke Torre-Healy, Jeffery M. Vahrenkamp, Kristofer C. Berrett, Hannah Conway, Chris Stubben, Richard Moffitt, Jason Gertz, Eric L. Snyder

**Affiliations:** Department of Pathology and Huntsman Cancer Institute, University of Utah, Salt Lake City, UT 84112, USA; Department of Biomedical Informatics & Stony Brook Cancer Center, Stony Brook University, Stony Brook, NY 11794; Medical Scientist Training Program, Stony Brook University, Stony Brook, NY 11794; Department of Oncological Sciences and Huntsman Cancer Institute, University of Utah, Salt Lake City, UT 84112, USA; Bioinformatics Shared Resource, Huntsman Cancer Institute, University of Utah, Salt Lake City, UT 84112, USA

**Keywords:** Pancreatic ductal adenocarcinoma, molecular subtype, HNF4α, SIX1, SIX4

## Abstract

Pancreatic ductal adenocarcinoma (PDAC) is an aggressive malignancy with a five-year survival of less than 5%. Transcriptomic analysis has identified two clinically relevant molecular subtypes of PDAC: Classical and Basal-like. The Classical subtype is characterized by a more favorable prognosis and better response to chemotherapy than the Basal-like subtype. The Classical subtype also expresses higher levels of lineage specifiers that regulate endodermal differentiation, including the nuclear receptor HNF4α. Using *in vitro* and *in vivo* PDAC models, we show that HNF4α restrains tumor growth and drives tumor cells toward an epithelial identity. Gene expression analysis from murine models and human tumors shows that HNF4α activates expression of genes associated with the Classical subtype. Although HNF4α loss is not sufficient for complete conversion to the Basal-like subtype gene expression profile, HNF4α directly represses SIX4 and SIX1, mesodermal lineage specifiers expressed in the Basal-like subtype. Finally, HNF4α-negative PDAC cells rely on expression of SIX4 and SIX1 for proliferation *in vitro* and *in vivo*. Overall, our data show that HNF4α regulates the growth and molecular subtype of PDAC by multiple mechanisms, including activation of the Classical gene expression program and repression of SIX4 and SIX1, which may represent novel dependencies of the Basal-like subtype.

## Introduction

Pancreatic ductal adenocarcinoma (PDAC) is an extremely aggressive malignancy with a five-year survival of less than 5% (1). Several recent genome-wide analyses of PDAC have identified discrete molecular subtypes of this disease (2–4). Most recently, the Cancer Genome Atlas Research Network reported an integrated genomic, transcriptomic and proteomic analysis of an additional 150 PDAC samples (5). This integrative analysis found significant overlap between the different subtypes, ultimately suggesting that PDAC can fundamentally be classified into two major subtypes based on cancer cell autonomous properties: Classical and Basal-like. These subtypes appear to be clinically relevant because the Classical subtype is associated with a better overall survival and better response to first line chemotherapy than the Basal-like subtype (2,4,6).

One major difference between these two subtypes is the relative levels of endodermal lineage specifiers, i.e. transcription factors that regulate development and differentiation of endodermally-derived tissues. The Classical subtype expresses high levels of these genes, including *HNF4A, GATA6*, *FOXA2*, *FOXA3*, and others (2). Histopathologic and gene expression analysis have shown that pancreatic intraepithelial neoplasia (PanIN, the most common non-invasive precursor lesion of PDAC) and well-differentiated PDACs exhibit a cellular identity that is distinct from the normal pancreas and is characterized by upregulation of transcripts highly expressed in the foregut. This suggests pancreatic neoplasia initially adopts a cellular identity resembling the tissue from which the pancreas originated during development (7, 8). Moreover, these correlations raise the question of whether specific endodermal lineage specifiers are not merely markers of discrete subtypes, but might directly regulate molecular subtype, malignant potential and therapeutic response in PDAC.

The nuclear receptor superfamily member HNF4α is a master regulator of epithelial differentiation in multiple tissues, including the gastrointestinal (GI) tract and the liver (9–12). HNF4α also regulates a variety of biological processes that impact tumor progression, including proliferation (13), metabolism (14), and inflammation (15). Consistent with genomics studies mentioned above, HNF4α protein is detectable in human pancreatic neoplasia, and its levels correlate with tumor differentiation state (16). Immunohistochemistry (IHC) on tissue sections has shown that PanIN lesions and well-differentiated PDAC express higher levels of HNF4α than the normal pancreas, whereas HNF4α is downregulated in poorly differentiated PDAC. Nine proteins that are part of an HNF4α-regulated network were identified by proteomic analysis of plasma from mice with PanIN lesions (16, 17), suggesting that HNF4α is functionally active in early pancreatic neoplasia.

In previous work, we have shown that HNF4α plays a critical role in a mouse model of invasive mucinous adenocarcinoma of the lung (18), a subtype of lung cancer that expresses many of the same foregut markers observed in the Classical subtype of PDAC. Based on these data and the differential expression of HNF4α between the Classical and Basal-like subtypes of human PDAC, we sought to determine whether HNF4α also plays a functional role in this disease. We found that HNF4α is a major activator of the Classical gene expression program and restrains PDAC growth in multiple models of the disease. Although HNF4α loss is not sufficient for complete subtype switching, HNF4α represses the expression of SIX4 and SIX1, two mesodermal lineage specifiers highly expressed in the Basal-like subtype of PDAC.

## Results

### *Hnf4a* deletion accelerates tumorigenesis in a mouse model of pancreatic ductal adenocarcinoma

To dissect the role of HNF4α in PDAC, we first used a mouse model of pancreatic neoplasia that closely mimics the human disease. This model employs the *Pdx1-Cre* transgene to activate expression of KRAS^G12D^ from its endogenous locus during pancreatic development, which thereby induces neoplasia that closely models human PanIN and PDAC (19, 20). Greater than 90% of human PanINs and PDACs contain mutations that result in constitutive activation of the *KRAS* oncogene, which is believed to be the initiating event in most cases (21). KRAS is a small GTPase that interacts with receptor tyrosine kinases at the plasma membrane to transduce growth factor-induced signals to several intracellular effectors (22).

*Pdx1-Cre; Kras^LSL-G12D^* mice develop acinar to ductal metaplasia (ADM) and murine PanIN (mPanIN) lesions with complete penetrance, and these *in situ* lesions occasionally progress to invasive cancer in older mice. To accelerate cancer progression, a conditional allele of the p53 tumor suppressor is incorporated into the model. Excision of one allele of *p53* (*Pdx1-Cre; Kras^LSL-G12D^*; *p53^F/+^* mice) accelerates neoplastic progression to invasive and metastatic adenocarcinomas, which often develop in mice 3-6 months of age (19). We initially evaluated a cohort of control mice (*Pdx1-Cre; Kras^LSL-G12D^*; *p53^F/+^*; *Hnf4a^+/+^*) at 12 weeks of age (n=17, Figure 1A-C). As expected from published reports (16), HNF4α was robustly expressed in preinvasive lesion (ADM and mPanIN) arising in these mice (Figure 1A, left). Careful evaluation of HNF4α levels in the subset of mice harboring PDAC (n=7) showed that 3/7 tumors were diffusely (>90%) HNF4α-positive. In contrast, one tumor contained a mix of HNF4α-positive and negative cells and 3/7 tumors were completely HNF4α-negative, despite the presence of adjacent HNF4α-positive mPanIN (Figure S1A). HNF4α-positive and mixed tumors exhibited a range of differentiation states (including well, moderately and poorly differentiated), whereas the three HNF4α-negative tumors were poorly differentiated. Taken together, these data shown that HNF4α is expressed at higher levels in early pancreatic neoplasia than normal pancreas, but that HNF4α can also be stochastically downregulated during tumor progression.

**Figure 1.**
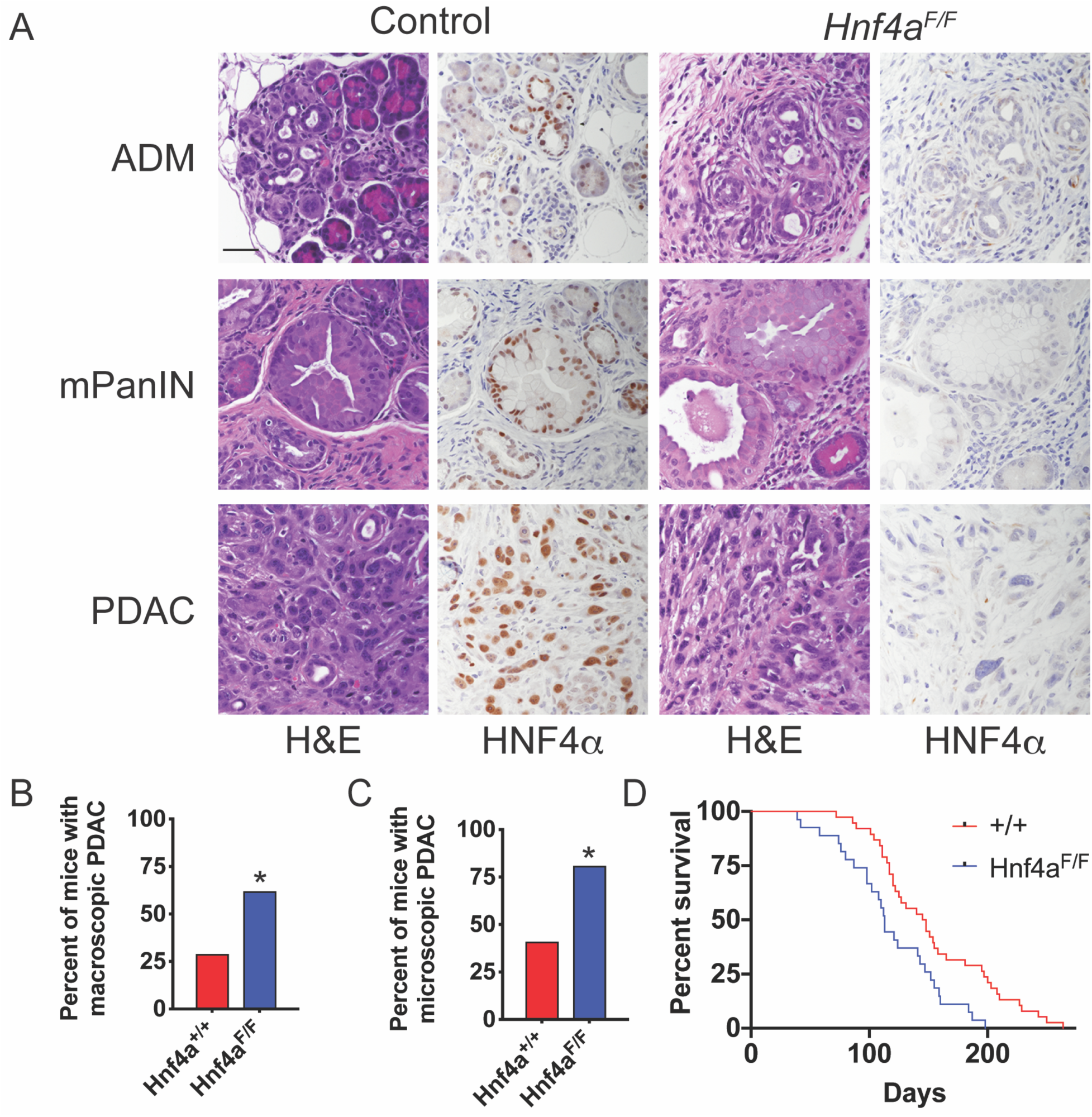
*Hnf4a* deletion accelerates tumorigenesis in a mouse model of pancreatic ductal adenocarcinoma. A. H&E and IHC for HNF4α on pancreatic neoplasia from *Kras^LSL-G12D/+;^ p53^F/+^; Pdx1-Cre; Hnf4a^F/^*^F^ mice and *Hnf4a^+/+^* controls at 12 weeks of age. Scale bar: 50 microns. ADM: acinar to ductal metaplasia. mPanIN: Pancreatic intraepithelial neoplasia. B-C. Percentage of *Kras^LSL-G12D/+;^ p53^F/+^; Pdx1-Cre; Hnf4a^F/F^* mice (n=21) and *Hnf4a^+/+^* controls (n=17) with macroscopic (B) and microscopic (C) PDAC at 12 weeks of age. *p<0.05, Chi-Square. D. Survival curve of *Kras^LSL-G12D/+;^ p53^F/+^; Pdx1-Cre; Hnf4a^F/^*^F^ mice (n=27) and *Hnf4a^+/+^* controls (n=38). p=0.0027, Log-rank.

Twelve isoforms of HNF4α have been identified to date (23), which arise as a result of transcription from two different promoters (P1 and P2) as well as alternative exon splicing. The P1 and P2 promoters are activated in a tissue-specific manner, and P1 and P2 isoforms have distinct roles in tumorigenesis (24, 25). Using monoclonal antibodies specific for the P1 and P2 isoforms, we found that HNF4α-positive neoplasia predominantly expressed the P2 isoform in this model (Figure S1B).

In contrast to control mice, all neoplasia (ADM, mPanIN and PDAC) in *Hnf4a^F/F^* mice at 12 weeks was HNF4α-negative (n=21, Figure 1A, right). We found no evidence of incomplete recombinants, which can be observed in the Cre-Lox system when there is selection for retention of a conditional allele. Although *Hnf4a* deletion had no obvious effect on the morphology or quantity of preinvasive lesions, it significantly increased the proportion of mice with either macroscopic PDAC (Figure 1B) or microscopic PDAC (Figure 1C) at 12 weeks of age.

In a separate survival analysis, we found that *Hnf4a* deletion significantly reduces survival in this model (median survival 113 days in *Hnf4a^F/F^* mice vs. 147 days in *Hnf4a^+/+^* mice, Figure 1D). Consistent with our analysis of tumors in 12-week old mice, a subset of tumors in *Hnf4a^+/+^* mice exhibited stochastic HNF4α downregulation, whereas tumors in *Hnf4a^F/F^* mice were consistently HNF4α-negative. Specifically, 9/20 tumors available for evaluation in *Hnf4a^+/+^* mice were HNF4α-mixed and 2/20 were HNF4α-negative (all of which were included in the overall survival anlaysis). Most tumors in the survival analysis harbored both moderately and poorly differentiated components, although a minority were entirely poorly differentiated (2/20 in *Hnf4a^+/+^*mice and 4/12 evaluated in *Hnf4a^F/F^* mice). Almost all survival mice (18/20 *Hnf4a^+/+^* and 11/12 *Hnf4a^F/F^* mice) harbored metastases to sites such as liver, kidney, peritoneal lymph nodes, diaphragm and the lung.

In the absence of p53 mutations, *Pdx1-Cre; Kras^LSL-G12D^* mice develop ADM and PanIN with high penetrance, but these lesions rarely progress to PDAC (19). To determine whether *Hnf4a* deletion would be sufficient for PDAC development, we aged a cohort of *Pdx1-Cre; Kras^LSL-G12D^; Hnf4a^F/F^*mice (and *Hnf4a^+/+^* controls, n=4-8 mice/group at each timepoint) and analyzed their pancreata at 6 and 9 months of age. We observed ADM and PanIN in nearly all mice of both genotypes, but no evidence of PDAC, despite the lack of HNF4α expression in *Hnf4a^F/F^* mice (Figure S1C).

Taken together, these data show that HNF4α restrains the growth of pancreatic neoplasia in this model system, predominantly at the stage of invasive PDAC. These results are also consistent with the biology of human PDAC, specifically the observation that Basal-like (HNF4α-low) tumors confer a worse prognosis, but that both HNF4α-positive and HNF4α-negative disease are highly lethal (2,4,6).

### HNF4α reconstitution impairs PDAC growth and imposes epithelial differentiation *in vitro* **and *in vivo***

To more precisely define the role of HNF4α in PDAC, we developed additional systems to manipulate its expression in established neoplasia. First, we generated two cell lines (HC800 and HC569) from PDAC arising in *Pdx1-Cre; Kras^LSL-G12D^*; *p53^F/+^*; *Hnf4a^F/F^* mice using standard two-dimensional (2D) culture conditions. We then stably transduced these cell lines with lentivirus enabling doxycycline (dox) induction of the HNF4α8 isoform, a major product of the P2 promoter. We used a P2 isoform for reconstitution studies because only HNF4α-P2 is expressed in the autochthonous model (Figure S1B). We then obtained single cell clones that exhibited optimal dox induction of HA-tagged HNF4α (Figure S2A) and subjected them to proliferation analysis. Exogenous HNF4α significantly inhibited the proliferation of both cell lines *in vitro* (Figure 2A) with no obvious induction of apoptosis as assessed by cleaved caspase-3 (Figure S2B). HC800 cells expressing exogenous HNF4α exhibited a more epithelial morphology *in vitro* (Figure S2C), suggesting that HNF4α reconstitution also modulated their differentiation state.

**Figure 2.**
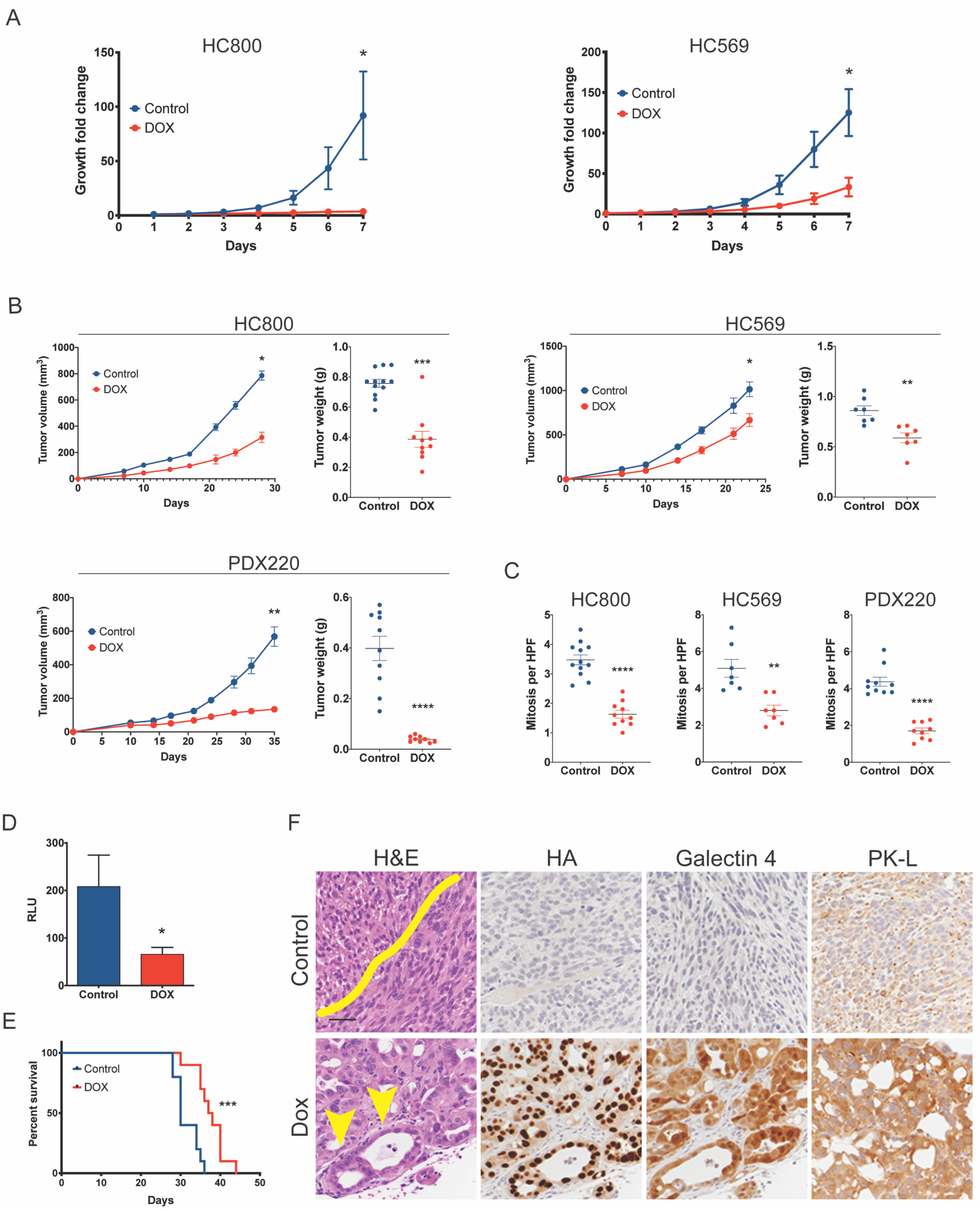
Restoration of HNF4α expression inhibits proliferation and induces epithelial differentiation (mesenchymal to epithelial transition) in PDAC. A. Quantitation of proliferation in murine (HC800 and HC569) PDAC cells stably expressing TRE-HNF4α8 cells the presence or absence of doxycycline (n=3 biological replicates, 2-3 independent experiments). Primary murine PDAC cell lines were single cell cloned. Graph represents mean +/-SEM. p= 0.03, Wilcoxon. B-C. HC800, HC569, and PDX220 cells stably transduced with TRE-HNF4α8 were injected subcutaneously into NOD-SCID-gamma chain deficient (NSG) mice. Mice were fed with doxycycline chow (red) or regular chow (blue) starting 1 week before transplantation. B. Flank tumor volume and mass are shown. C. Flank tumor growth was assessed by measuring mitoses per high power field (HPF). Data represented as mean ± SEM. *p< 0.05; **p< 0.01; ***p< 0.001 by Wilcoxon test (for tumor volume) or Mann-Whitney test (for tumor mass and mitoses). D-F. HC800-TRE-HNF4α8 cells were injected intraperitoneally into NSG mice. Mice received either doxycycline chow or control chow starting 1 week prior to cell injection (n=10 mice per group). Tumor cells were stably transduced with luciferase expression vector prior to injection. D. Quantitation of relative light units (RLU) in each group 20 days post flank injections. Graph shows fold change in RLU normalized to basal measurements. Data represented as mean ± SEM. *p<0.05 by t test. E. Survival analysis. ***p<0.001 by log-rank test. F. Representative H&E and IHC for HA-tagged HNF4α and target genes in intraperitoneal tumors from each group. Yellow line demarcates two fascicles of tumor cells oriented in opposite directions. Yellow arrows mark glandular differentiation. Scale bar: 100 microns.

To test the effect of HNF4α restoration *in vivo*, we first injected cells subcutaneously into mice fed control or dox-containing chow to induce HNF4α. HNF4α was robustly expressed in tumors of mice on dox chow (Figure S2D). Exogenous HNF4α significantly inhibited tumor growth of both murine PDAC cell lines as assessed by both volume and mass (Figure 2B), and this was accompanied by a significantly lower proliferation rate than controls (Figure 2C). To better recapitulate the biology of PDAC, we also evaluated the growth of HC800 cells in the peritoneal space, which is one of the first sites of extra-pancreatic dissemination as PDAC progresses. We found that these cells grew readily in the peritoneum, forming numerous macroscopic nodules on the serosal surface of intraperitoneal organs. In a formal survival study, dox-mediated induction of HNF4α reduced tumor burden at 20 days post injection (Figure 2D) and significantly increased overall survival (Figure 2E).

In addition to restraining proliferation, HNF4α had a clear impact on differentiation state *in vivo*, shifting tumors toward a more well differentiated epithelial morphology (Figures 2F and S2D). This was particularly striking in the HC800 cell line. Control HC800 tumors had a uniform morphology composed of spindled cells growing in a fascicular pattern that is characteristic of a sarcomatoid or quasi-mesenchymal differentiation state (Figure 2F, top row). IHC for the HA tag confirmed that exogenous HNF4α was not expressed in these tumors. In contrast, tumors growing in mice fed dox chow adopted the morphology of a moderately to poorly differentiated adenocarcinoma, with a glandular rather than fascicular growth pattern and cells that were round rather than spindled. These tumors expressed uniformly high levels of HA-tagged HNF4α, its target PK-L, and Galectin 4, a marker of foregut differentiation (Figure 2F, bottom row).

To further evaluate these results in human PDAC cells, we stably transduced a human PDAC cell line derived from the xenograft PDX220 (26) with lentivirus encoding dox-inducible HNF4α8 and tested the effect of HNF4a induction on subcutaneous tumor growth. Control PDX 220 tumors contain a mixture of HNF4α-positive and -negative cells (Figure S2E). Interestingly, HNF4α-positive cells are a moderately differentiated adenocarcinoma, whereas the HNF4α-negative cells consist of squamous cell carcinoma that expresses *Δ*Np63, a marker of squamous differentiation (Figure S2E-F). Exogenous HNF4α substantially inhibited overall PDX220 tumor growth and cellular proliferation (Figure 2B-C). Moreover, tumors arising in dox-treated mice consisted entirely of adenocarcinoma that was better differentiated than control tumors (Figure S2F), showing that HNF4α not only suppressed squamous differentiation in this model but also modulated the differentiation state of the adenocarcinoma component.

Taken together, these results show that restoring HNF4α function can induce major changes in differentiation state and inhibit PDAC growth both *in vitro* and *in vivo*.

### *Hnf4a* deletion in PDAC organoids alters differentiation and three dimensional growth pattern *in vivo*

Given that HNF4α is often stochastically downregulated during PDAC progression, we next developed a system that would enable us to inactivate endogenous *Hnf4a* in established murine pancreatic neoplasia. In mouse models of lung cancer, we have previously used a sequential recombinase strategy to delete genes in established tumors (18). Merging this approach with pancreatic organoid culture systems (27), we established PDAC organoids by enzymatically dissociating pancreata from *Kras^FSF-G12D/+;^ p53^Frt/Frt^; Rosa26^FSF-CreERT2^; Hnf4a^F/F^* mice. We then treated these cells *in vitro* with adenovirus expressing the FlpO recombinase, which activated transcription of oncogenic KRAS^G12D^ and inactivated p53, leading to the emergence of HNF4α-positive PDAC organoids (Figure 3A and S3A). FlpO also activated transcription of Cre^ERT2^ from the *Rosa26* locus. This enabled us to treat these cultures with 4-hydroxy-tamoxifen (4-OHT), thereby driving Cre^ERT2^ into the nucleus where it excised the respective conditional alleles. IHC and qRT-PCR demonstrated loss of HNF4α as well as its target *Pklr* in four independently derived organoid cultures (Figure S3A-B).

**Figure 3.**
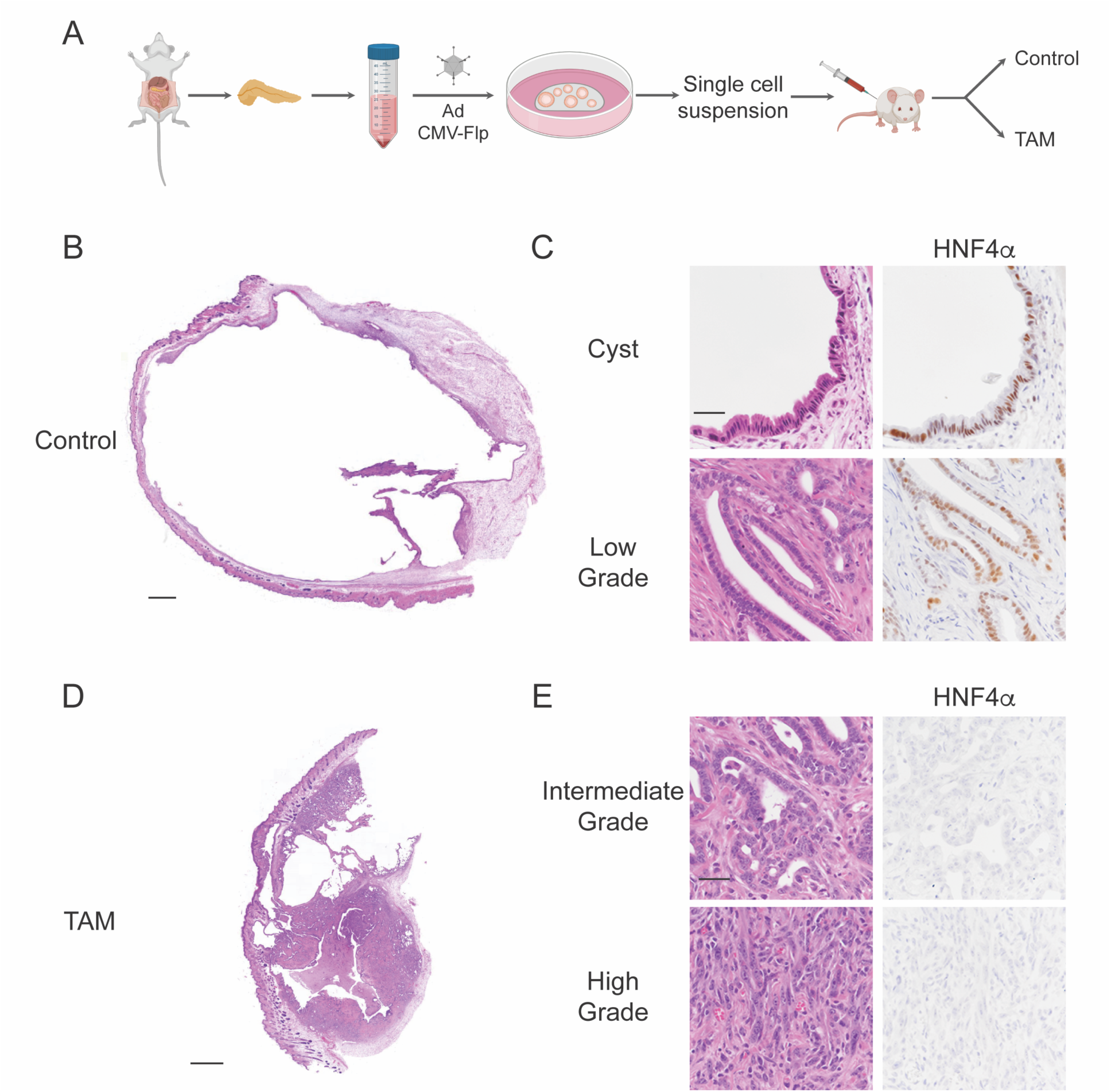
*Hnf4a* deletion alters differentiation and three dimensional growth in murine organoid models of PDAC. A. Schematic of organoid derivation from pancreata of *Kras^FSF-G12D/+;^ p53^Frt/Frt^; Rosa26^FSF-CreERT2^; Hnf4a^F/^*^F^ mice. B-E. H&E and IHC analysis of a single cell suspension of SC1853 organoids injected subcutaneously into NSG mice. Mice were fed control (B-C) or tamoxifen (D-E) chow starting at 1 week prior to organoid implantation. Tumors were analyzed at 6 weeks post injection. B and D: scanning magnification, scale bar: 1 mm. C and E: High power images, scale bar: 100 microns.

We next asked whether loss of HNF4α would affect the growth and differentiation state of these organoid lines *in vivo*. We injected two lines (SC1853 and SC1693) subcutaneously into NOD/SCID-gamma chain deficient (NSG) mice. In mice treated with control chow, both lines formed macroscopic, fluid filled cysts (Figure 3B and S3C). These cysts were predominantly lined by a single layer of HNF4α-positive columnar epithelial cells (Figure 3C and S3D, top row). We also identified small areas of well-differentiated adenocarcinoma (Figure 3C and S3D, bottom row) within the walls of these cystic structures.

In contrast, tumors that grew in mice on tamoxifen chow exhibited a completely distinct three-dimensional growth pattern and differentiation state. These tumors consisted of a mixture of invasive adenocarcinoma and microscopic cystic structures (Figure 3D and S3E). Moreover, the adenocarcinoma component was less well differentiated than control tumors. *Hnf4a*-deleted tumors from both organoid lines exhibited areas of moderately differentiated (intermediate grade) HNF4α-negative adenocarcinoma (Figure 3E and S3F). Moreover, *Hnf4a* deletion in tumors of the SC1853 organoid was sufficient to induce areas of high grade, poorly differentiated adenocarcinoma (Figure 3E, bottom row).

### HNF4α activates the Classical gene expression program in PDAC

Given its impact on the growth and differentiation state in multiple models of PDAC, we sought to formally test the hypothesis that HNF4α regulates molecular subtype in PDAC. As expected from prior studies (2, 5), we observed a robust correlation (r = 0.71, p = 9.1e-13) between *HNF4A* expression and Classical Score in high purity tumors from the TCGA-PAAD data (Figure 4A, left). In comparison, *HNF4A* levels and Basal-like score were weakly negatively correlated (r = −0.48, p = 1.4e-5) in this dataset (Figure 4A, right). We also evaluated the relationship between *HNF4A* levels and survival in high purity tumors (>30% cellularity) from two independent datasets (2, 5) by either splitting cases around median HNF4A expression or into upper and lower quartiles of *HNF4A* expression (Figure S4A). In the PACA-AU dataset, patients with tumors in the upper quartile of *HNF4A* expression survived significantly longer than in the lowest quartile (p=0.012), consistent with known survival differences between the Classical and Basal-like subtypes. Other analyses showed a similar (though non-significant) trend. Interestingly, in the TCGA dataset, a subset of patients with *HNF4A*-high tumors were still alive at the time of analysis, whereas none of the patients in the *HNF4A*-low cohort survived beyond the last recorded time point (2000 days).

**Figure 4.**
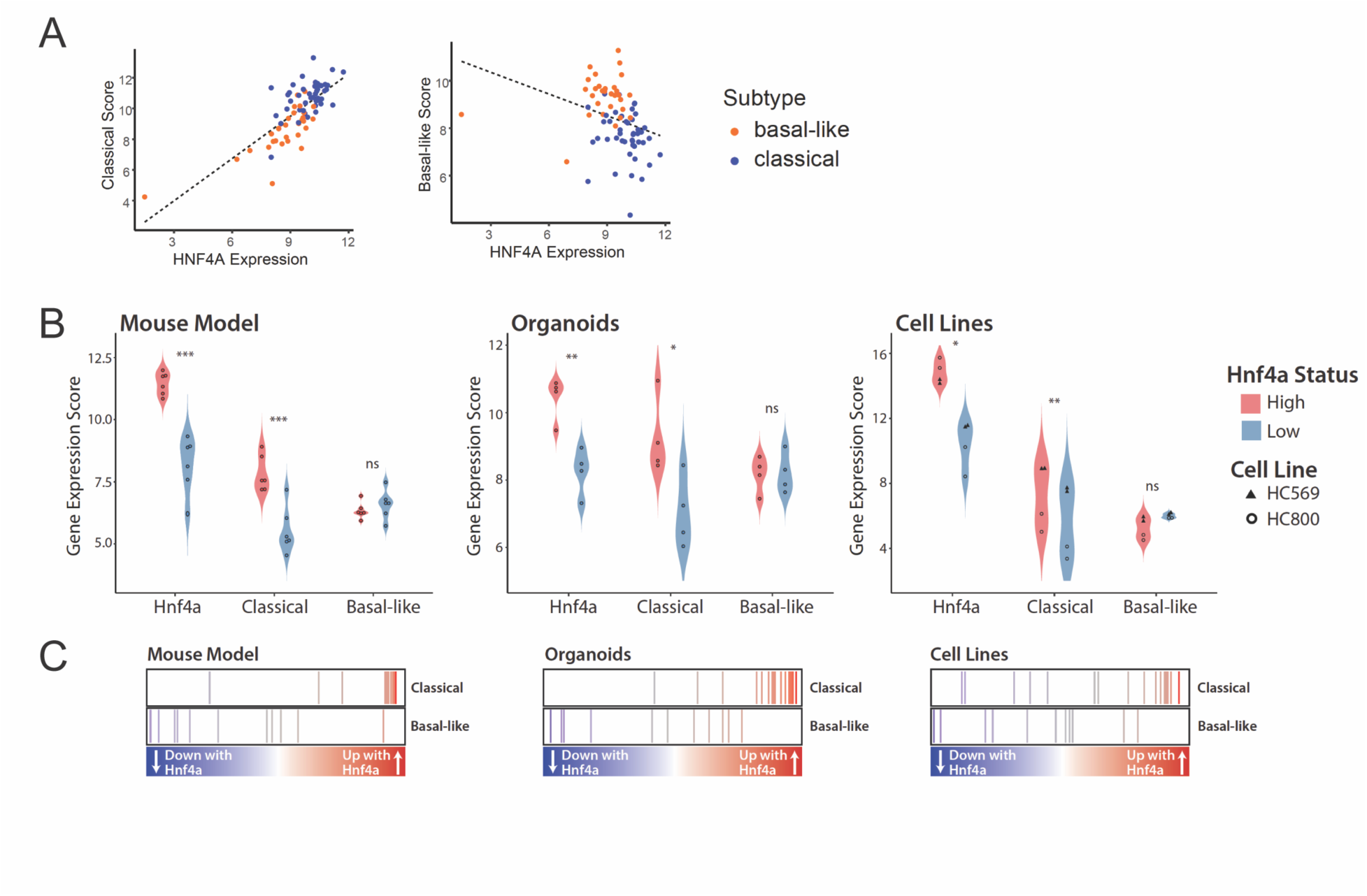
HNF4α regulates PDAC molecular subtype. A. Correlation between HNF4A expression in patients with Classical Score (r = 0.71, p= 9.1e-13) and Basal-like score (r = −0.48, p= 1.4e-5) in TCGA-PAAD data. B. Scores were generated for Basal-like and Classical subtype expression from RNA-seq data. Each point represents one sample, and samples were grouped by their HNF4*a* status. Subtype scores were calculated using mouse homologs of the established Moffitt et al. signatures. p-values obtained via t-test (paired for cell lines). ***p<0.001, **p<0.01, *p<0.05. C. Gene Set Enrichment Analysis was performed comparing the HNF4*a*-positive samples to the HNF4*a*-negative samples, and their respective Signal-to-Noise Ratios (SNR) are plotted. Red indicates that expression of the gene increases with HNF4*a* expression, while blue indicates expression decreases.

Next we performed gene expression analysis on autochthonous murine PDAC tumors from *Hnf4a^+/+^* and *Hnf4a^F/F^*mice (n=6 per genotype), using IHC to exclude any tumors from wild type mice that had stochastically downregulated HNF4α. We identified a total of 597 differentially expressed genes in this analysis (Supplemental Table 1), including known HNF4α targets in normal tissue such as *Pklr* and *Apob*. To determine whether *Hnf4a* deletion altered molecular subtype of these tumors, we assigned a Classical and Basal-like score to each dataset based on mouse homologues of the scores described in (4). The Classical score of *Hnf4a*-deleted tumors was significantly lower than *Hnf4a^+/+^* tumors (Figure 4B-C, left panels). In contrast, the Basal-like score was not significantly different between the two groups, although there was a slight trend toward a higher score in the *Hnf4a*-deleted tumors. Evaluation of lineage specifiers that regulate endodermal differentiation (Figure S4B) showed that some appear to be HNF4α targets in this model (e.g., HNF1α, FoxA3 and CDX2), whereas others are not significantly affected by *Hnf4a* deletion, suggesting they are situated adjacent to or upstream of HNF4α in the hierarchy of lineage specifiers specifying the Classical subtype of PDAC.

Using Illumina Correlation Engine, we performed pathway analysis on genes differentially expressed between HNF4α-positive and HNF4α-negative autochthonous tumors (Supplemental Table 2). GO terms and pathways associated with epithelial differentiation (e.g., “cell-cell adhesion”) and other known functions of HNF4α (e.g., “Maturity onset diabetes of the young” and various metabolic pathways) were significantly enriched in HNF4α-positive tumors. In contrast, GO terms associated with mesodermal differentiation (e.g., “regulation of neurogenesis” and “skeletal system development”) were significantly enriched in HNF4α-negative tumors. Consistent with these results, the Body Atlas analysis tool showed that the gene expression profile of HNF4α-positive tumors correlates highly with endodermally derived tissues, such as the GI tract, whereas HNF4α-negative tumors correlate with mesodermally derived tissues found in the central nervous system and cardiovascular system (Figure S4C and Supplemental Table 2).

To expand upon these observations and determine whether HNF4α can regulate PDAC identity *in vitro* in a cell-autonomous manner, we performed gene expression analysis on the previously described cell lines and organoids isogenic for HNF4α expression (Supplemental Tables 3 and 4). In both cell line systems, HNF4α-positive cells had a significantly higher Classical score than HNF4α-negative cells (Figure 4B-C, middle and right hand panels). In contrast, the Basal-like score did not significantly change with HNF4α-expression. Intersection of all three datasets identified a total of 49 genes that were upregulated by HNF4α in all model systems (Figure S4D and Supplemental Table 5). Although few genes were downregulated by HNF4α in all model systems, primarily due to lack of overlap with the organoid dataset, we identified 65 genes downregulated by HNF4α both in vivo and in 2D cell lines (Figure S4D and Supplemental Table 5). Taken together, these data show that HNF4α is a critical regulator of cellular identity and molecular subtype in multiple models of PDAC. Specifically, HNF4α is required for activation of the complete Classical gene expression program. Although loss of HNF4α is not sufficient for full conversion to a Basal-like state, a subset of genes associated with the Basal-like subtype appear to be repressed by HNF4α.

### Analysis of HNF4α-chromatin interactions in PDAC

In order to identify direct HNF4α target genes and investigate the impact of HNF4α on chromatin accessibility, we performed ChIP-seq with an antibody that recognizes the HA-tagged HNF4α as well as ATAC-Seq in HC800 and HC569 cells. For both assays, cells were treated with dox or control media for one week prior to chromatin analysis. We identified 18,419 and 15,593 reproducibly significant peaks by HA ChIP-seq in HC569 and HC800 dox-treated cells, respectively. We found that 84.4% of all sites were present in both cell lines, showing a strong degree of concordance in HNF4α binding (Figure 5A and S5A). As expected, motifs bound by HNF4α and HNF4*γ* were the most significantly enriched in ChIP-seq peaks (Figure S5B). Examination of HNF4α binding distribution showed that 45% of peaks localized to promoters, whereas 13% were found in distal regions (Figure 5B). We corroborated HNF4α occupancy in relevant and well-known direct target genes such as *Pklr, Lgals4*, and *Hnf1a* (Figure 5C). To explore the connection between binding sites and gene expression, we intersected our ChIP-seq and RNA-seq datasets. We observed 69% of the differentially expressed genes displaying binding sites for HNF4α (Figure 5D). For those genes bound by HNF4α, we found that 61% were upregulated and 39% downregulated, indicating that HNF4α preferentially acts as a transcriptional activator in this context.

**Figure 5.**
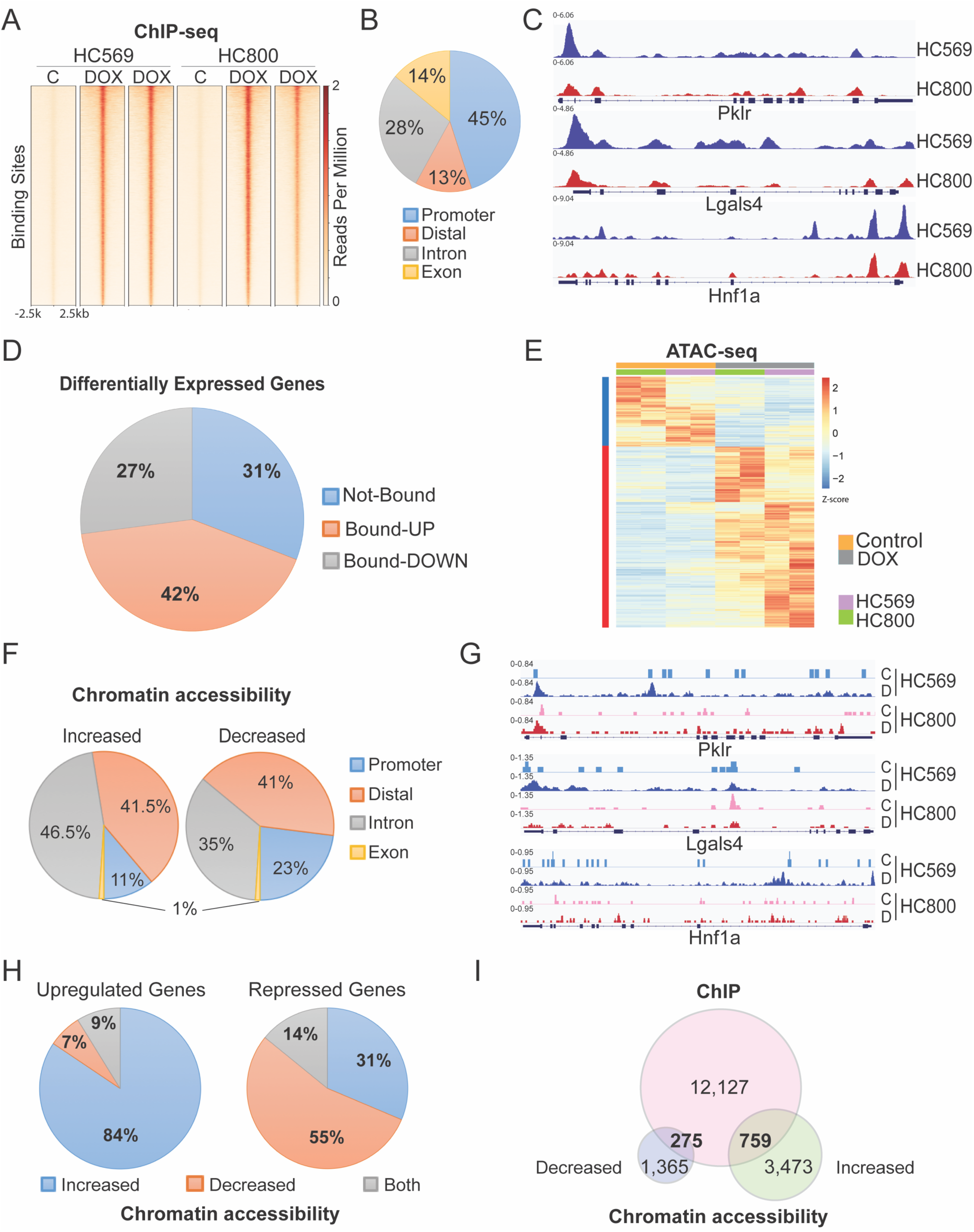
Global analysis of HNF4α occupancy and chromatin accessibility landscape in PDAC cells. A. Heatmaps displaying HNF4α binding sites in a 5-kb window around the ChIP-seq peak summits in HC569 and HC800 cells. B. Pie chart displays the genomic distribution of HNF4**α** binding sites. C. Representative browser track images of three known direct targets of HNF4α in HC569 (blue) and HC800 (red). D. Overlap between RNA-seq and ChIP-seq data. Distribution of differentially expressed genes bound and not bound by HNF4α. Bound-UP: upregulated genes with HNF4α occupancy. Bound-DOWN: downregulated genes bound by HNF4α. E. Heatmap shows signal at ATAC-seq regions that increase (red line) and decrease (blue line) chromatin accessibility upon dox-mediated HNF4α induction in HC800 (green) and HC569 (purple) cell lines. F. Distribution of genomic features for ATAC-seq regions with increased and reduced chromatin accessibility. G. Representative browser track images showing intensity of ATAC-seq signal at the indicated genes in dox-treated cells (blue: HC569, red: HC800) compared to controls (light blue: HC569, pink: HC800) H. Overlap between RNA-seq and ATAC-seq data sets. Distribution of up-and downregulated genes between more and less accessible chromatin. I. Venn diagram displays the overlap between HNF4α-bound sites and ATAC-seq peaks with increased or reduced genomic accessibility.

We further assessed the effect of HNF4α on global chromatin accessibility by performing ATAC-seq in both cell lines upon 1 week treatment with doxycycline or vehicle. Principal component analysis showed that control (HNF4α-negative) HC569 and HC800 cells clustered separately, but that HNF4α restoration caused similar changes in chromatin accessibility in both cell lines (Figure S5C). A total of 5,872 significant ATAC-seq peaks were identified, with 4,232 sites exhibiting increased accessibility and 1,640 sites exhibiting decreased accessibility in the presence of HNF4α (Figure 5E). Genomic distribution analysis of ATAC-seq peaks showed that most of the sites with significant changes in accessibility are found in distal regions and introns. Despite not being the most frequent genomic feature found in differentially accessible chromatin, promoters were overrepresented in closed genomic sites relative to chromatin with increased accessibility (Figure 5F). We observed increased signal of ATAC-seq peaks in HNF4α target genes that are transcriptionally activated in dox-treated samples (Figure 5G). On further inspection, we observed that genomic regions with increased accessibility were enriched in motifs for binding of AP1 and HNF4α, among other nuclear receptor family members. In contrast, while closed chromatin peaks lack HNF4α binding motifs, they show significant enrichment for AP1 binding sites as well (Figure S5D).

Intersection of ATAC-seq and RNA-seq data identified 743 differentially expressed genes (488 upregulated and 255 silenced genes) that displayed changes in chromatin accessibility upon HNF4α restoration. The vast majority of the upregulated genes correlated with genomic regions with increased accessibility (84%), while most of the repressed genes were associated with ATAC-seq peaks that had reduced genomic accessibility (55%), although a considerable percentage (31%) correlated with genomic areas that become more accessible (Figure 5H and S5E).

Finally, correlation of HNF4α binding sites with ATAC-seq data demonstrated that less than 10% of the HNF4α bound sites displayed changes in chromatin accessibility and less than 20% of the ATAC-seq peaks with either more or less accessibility were occupied by HNF4α, suggesting that other transcription factors may be contributing to the HNF4α-dependent alterations in chromatin accessibility (Figure 5I and S5F).

Altogether, these results demonstrate that most of the differentially expressed genes that define the transcriptional network regulated by HNF4α in PDAC constitute direct targets of this endodermal cell fate determinant. Additionally, our data suggest that most of the changes in chromatin accessibility seem to be indirect effects of HNF4α restoration and not caused by its direct binding.

### HNF4α inhibits the expression of mesodermal lineage specifiers SIX4 and SIX1 in PDAC

Next, we sought to identify mechanisms by which loss of HNF4α might lead to de-repression of a subset of genes associated with the Basal-like subtype. *Six4*, a member of the *sine oculis* homeobox family, is the most differentially expressed transcription factor in *Hnf4a*-deleted autochthonous tumors compared with controls (Figure 6A). Moreover, *SIX4* expression is positively correlated with the Basal-like subtype of human PDAC and negatively correlated with Classical Score and *HNF4A* expression (Figure S6A). The *Six4* paralogue *Six1* is also differentially expressed in *Hnf4a*-deleted autochthonous tumors. Although SIX1 levels do not correlate with PDAC subtype to the same degree as SIX4 (Figure S6B), SIX1 levels positively correlate with SIX4 in human PDAC (Figure S6C). Overall, it appears that a subset of SIX4-high human PDAC also express high levels of SIX1 (Figure S6C).

**Figure 6.**
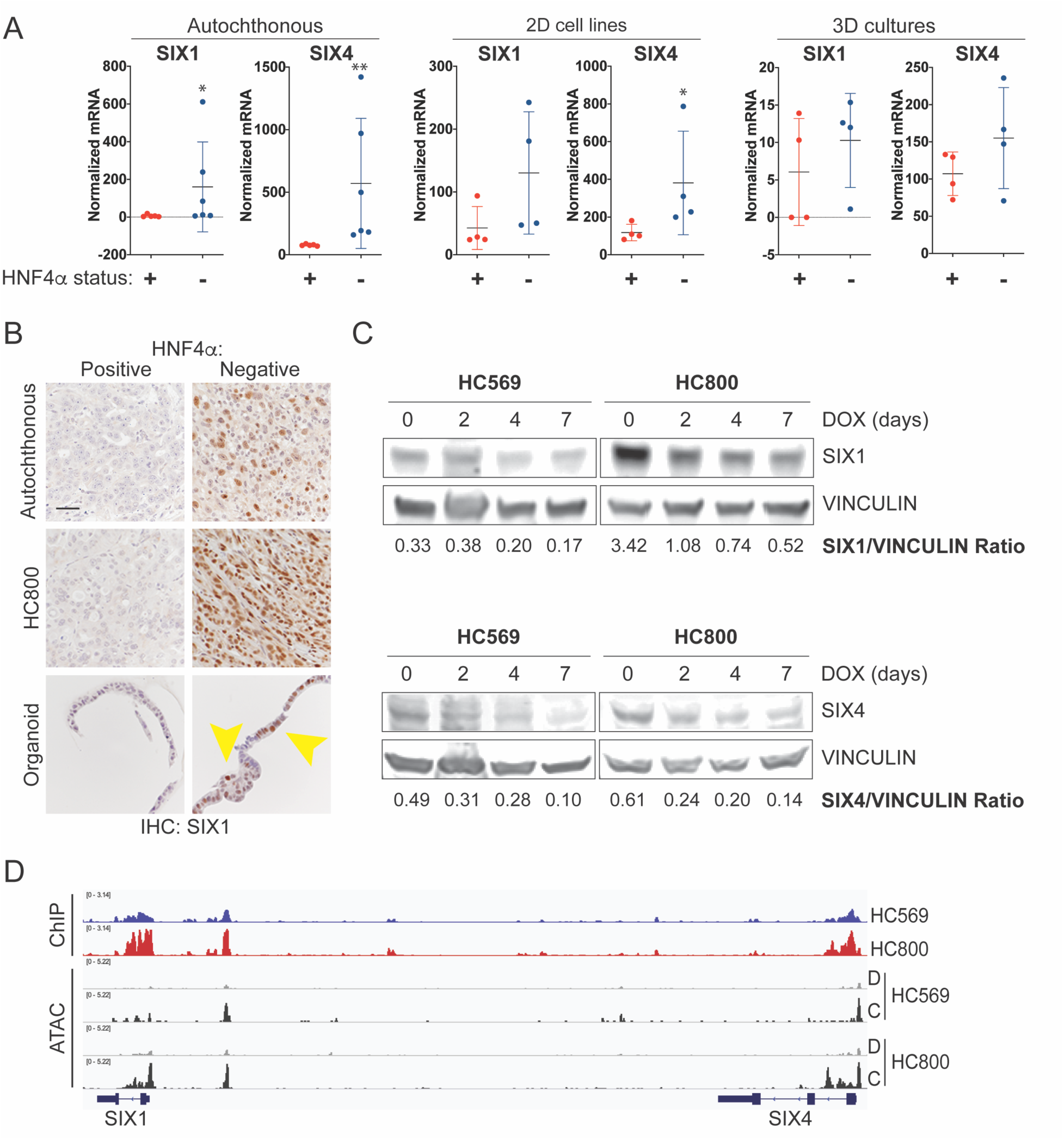
HNF4α inhibits the expression of mesodermal lineage specifiers SIX1 and SIX4. A. Six1 and Six4 mRNA levels in indicated samples as assessed by RNA-Seq. B. Representative IHC for SIX1 from indicated PDAC models with different HNF4α expression levels. Autochthonous: *Pdx1-Cre; Kras^LSL-G12D^*; *p53^F/+^* tumors from Hnf4a^+/+^ (left) or Hnf4a^F/F^ (right) mice. HC800: intraperitoneally injected cell line with dox-inducible HNF4α. Organoid: derived from *Kras^FSF-G12D/+;^ p53^Frt/Frt^; Rosa26^FSF-CreERT;^ Hnf4a^F/F^* mice. Three dimensional cultures were treated *in vitro* with vehicle control (left) or 4-hydroxy-tamoxifen (right) to delete *Hnf4a*. Yellow arrows indicate SIX1-positive cells. Scale bar: 100 microns. C. Representative immunoblot for SIX1 (50ug) and SIX4 (100ug) in doxycycline-treated murine PDAC cell lines. Ratio between SIX1 or SIX4 and Vinculin (loading control) intensities are shown. D. Browser track images display signal intensity of ChIP-Seq and ATAC-Seq in 2D lines. Blue: HNF4α ChIP in HC569. Red: HNF4α ChIP in HC800. Grey: ATAC peaks in dox-treated cells. Black: ATAC signal in control cells.

*SIX4* and *SIX1* are located immediately adjacent to each other in both mouse and human genomes and are co-expressed in several tissues during development (28). Both transcription factors bind the MEF3 DNA motif, and genetic knockout experiments have demonstrated their functional redundancy in the development of several mesodermally derived tissues, including skeletal muscle, sensory neurons, gonads and kidney (reviewed in (29)). SIX1 has been reported to promote the growth of the mesenchymal/Basal-like human PDAC cell lines Panc1 and MiaPaCa-2 (30, 31). However, there has been no systematic evaluation SIX4 in PDAC, either alone or in combination with SIX1, despite their co-expression in human tumors and evidence of significant functional redundancy during development.

To further investigate the ability of HNF4α to regulate SIX1 and SIX4 expression, we evaluated RNA and protein levels in all of our mouse models. Analysis of RNA-Seq data from individual tumors showed that *Six1* and *Six4* were not expressed in *Hnf4a^+/+^* autochthonous tumors (Figure 6A, left). Three out of six *Hnf4a^F/F^* tumors exhibited de-repression of both genes, whereas the other three *Hnf4a^F/F^* tumors had very low levels of *Six1* and *Six4* transcripts (similar to control tumors). We were able to evaluate SIX1 protein levels by IHC in an expanded panel of autochthonous tumors. We found that SIX1 was largely undetectable in *Hnf4a^+/+^* autochthonous tumors (Figure 6B, top row). In contrast, a panel of *Hnf4a^F/F^* tumors from 12-week old mice (n=14) exhibited a heterogenous pattern of SIX1 expression. SIX1 protein was detectable in 8 out of 14 *Hnf4a^F/F^*tumors, whereas the other 6 tumors were negative. Within the 8 positive tumors, the percentage of SIX1-positive cells ranged from less than 5% to greater than 75% (Figure 6B, top row and data not shown).

We next evaluated the ability of HNF4α to regulate SIX1 and SIX4 in our other model systems. Exogenous HNF4α significantly inhibited *Six1/4* mRNA expression in both *Hnf4a*-deleted cell lines (HC569 and HC800) (Figure 6A, middle panels), which led to decreased SIX1 and SIX4 protein levels *in vitro* and correlated with virtually undetectable SIX1 expression *in vivo* (Figure 6B, middle row, and 6C). RNA-Seq analysis of organoids showed a non-significant trend toward increased *Six4* levels after *Hnf4a* deletion (Figure 6A, right). Although *Six1* did not score as a differentially expressed gene in organoids, we were able to detect differences at the protein level. Control murine PDAC organoids were uniformly SIX1-negative, but SIX1 protein was detectable in two of the four *Hnf4a*-deleted organoid lines (SC1640 and SC1853, Figure 6B, lower row). ChIP-seq revealed that HNF4α directly binds both *Six1* and *Six4* genes in proximity to their promoter region in both cell lines. We also identified a significant peak in the intergenic region between *Six1* and *Six4*. Moreover, HNF4α occupancy in this region was accompanied by reduction in the chromatin accessibility (Figure 6D).

Taken together, these data show that HNF4α represses SIX1 and SIX4, two lineage specifiers that drive mesodermal development, likely through direct binding to their regulatory elements. However, loss of HNF4α expression is not sufficient for induction of SIX1 and SIX4 in all contexts, suggesting that other redundant mechanisms help repress SIX1/4 expression in the Classical subtype of PDAC.

### SIX1 and SIX4 are required for growth of HNF4α-negative PDAC *in vitro* and *in vivo*

Given that *Six* genes are frequently dysregulated in different cancers, we decided to further explore the oncogenic function and potential redundancy of SIX1 and SIX4 in HNF4α-negative PDAC. We initially performed CRISPR-Cas9 genome editing to ablate *Six1* and *Six4* in HC569 and HC800 cells by using 2 independent single guide RNA sequences targeting *Six1* and 4 targeting *Six4*, which were validated by immunoblotting (Figure S7A). We used the three most efficient sgRNAs targeting *Six4* for conducting functional studies.

We first assessed the impact of SIX1 deletion in cell proliferation of the bulk population of both murine PDAC cell lines *in vitro.* HC569 cells stably transduced with either sgRNA against SIX1 showed a significant reduction in cell proliferation, indicating this cell line is susceptible to *Six1* loss (Figure 7A). In contrast, loss of SIX1 expression in HC800 cells caused little to no change in proliferation relative to controls (Figure S7B). Single cell clones derived from these bulk populations revealed a similar pattern: *in vitro* proliferation of HC569 clones was remarkably impaired whereas HC800 clones showed almost no change (Figure 7B and S7C). In order to further explore the effect of *Six1* loss *in vivo*, we injected single cell clones of each sgRNA into NSG mice. SIX1-negative clones formed tumors that were significantly smaller and less proliferative than controls (Figure 7C). IHC confirmed the absence of SIX1 expression in tumors arising from both *Six1*-deleted clones, whereas SIX1 was readily detectable in control tumors (Figure 7D).

**Figure 7.**
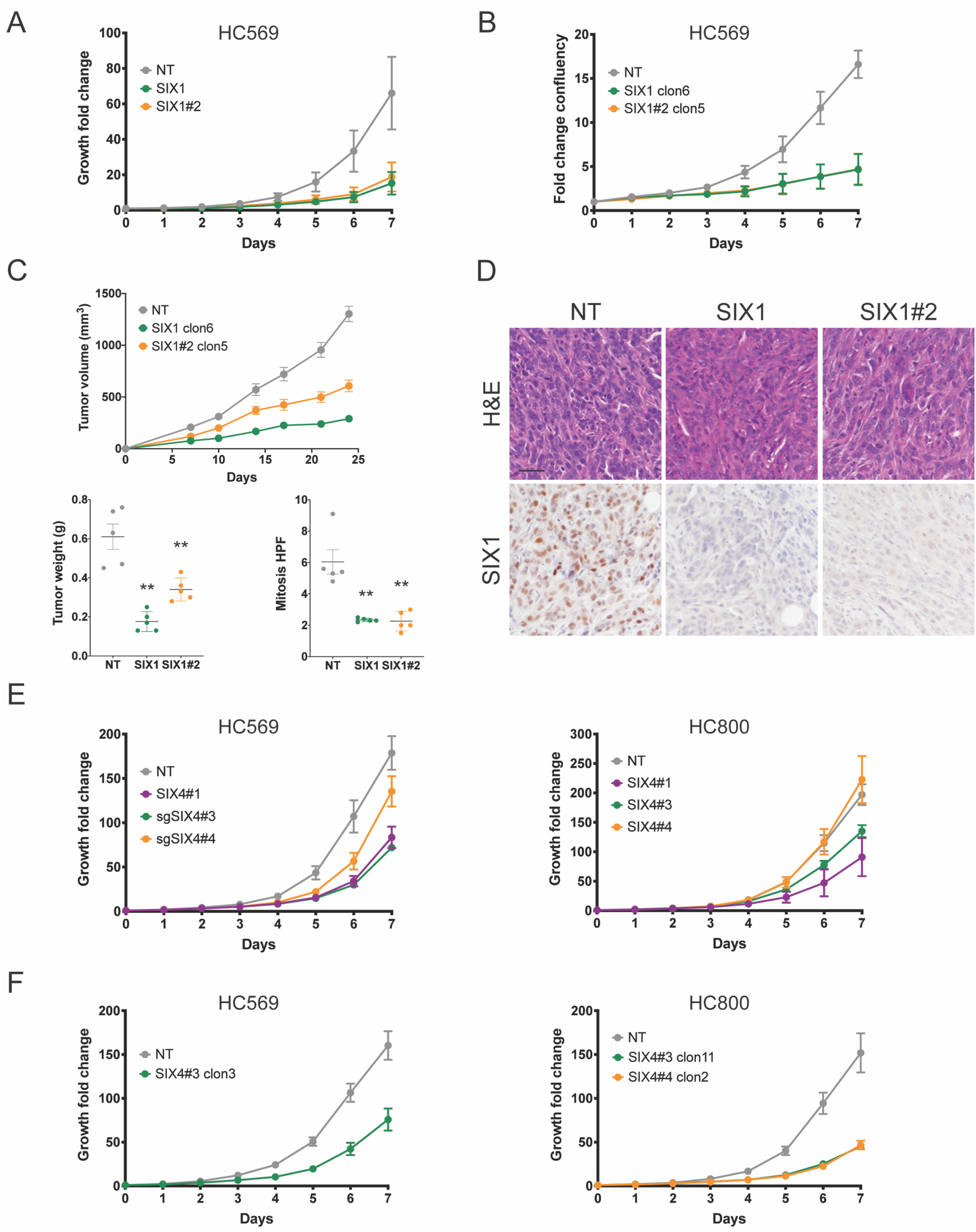
SIX4 and SIX1 are required for the growth of HNF4α-negative PDAC. A. Quantitation of proliferation in HC569 cells stably expressing Cas9 and the indicated sgRNA against *Six1* (n= 3 biological replicates, 2 independent experiments). Graph represents mean +/- SEM. p≤ 0.05, Wilcoxon test. B. Quantitation of proliferation in single cell cloned HC569 cells stably expressing Cas9 and the indicated sgRNA against *Six1* (n= 3 biological replicates, 2 independent experiments). Data represents mean +/- SEM. p≤ 0.05, Wilcoxon test. C-D. Individual clonal populations of *Six1* deleted-HC569 cell line were injected subcutaneously into the flank of NSG mice. Tumor volume was measured biweekly starting 1 week post implantation (n=5 mice per group). C. Flank tumor volume and weight, and mitosis per high power field are shown. Data represented as mean ± SEM. *p< 0.05; **p< 0.01; ***p< 0.001 by Wilcoxon test (for tumor growth) or Mann-Whitney test (for tumor weight and mitosis). D. Representative H&E and IHC for SIX1 are shown for each cohort. Scale bar: 100 microns. E. Quantitation of proliferation in HC800 and HC569 cells stably expressing Cas9 and the indicated sgRNA against *Six4* (n= 3 biological replicates, 2 independent experiments). Graph represents mean +/-SEM. p≤ 0.05, Wilcoxon test. F. Quantitation of proliferation in single cell cloned HC569 and HC800 cells stably expressing Cas9 and the indicated sgRNA targeting *Six4* (n= 3 biological replicates, 2 independent experiments). Graph represents mean +/- SEM. p≤ 0.05, Wilcoxon test.

*Six4* knock out caused a reduction in the growth rates of both cell lines compared with control cells. In particular, HC569 showed a significant reduction in proliferation with all three sgRNAs, while only sgRNAs SIX4#1 and #3 caused a modest decrease in HC800 growth capacity (Figure 7E). We derived single cell clones of HC569 and HC800 SIX4 knock out cell lines and observed that SIX4-deleted cells displayed a slower growth compared to their control counterparts (Figure 7F).

Taken together, these results show that proliferation of both HNF4α-negative PDAC cell lines tested is dependent on SIX4 expression. One of the two lines tested is also highly dependent on SIX1 for both *in vitro* proliferation and *in vivo* tumorigenesis, consistent with prior reports in human cell lines (30, 31).

## Discussion

Although two major molecular subtypes of human PDAC have been characterized clinically, the molecular regulators of their distinct biologic properties and clinical behaviors have not been fully identified. Here we show that the nuclear receptor HNF4α is a critical, non-redundant regulator of growth and molecular subtype in PDAC. Using complementary *in vitro* and *in vivo* model systems, we show that HNF4α restrains tumor growth and enforces an epithelial differentiation program by at least two distinct mechanisms. First, HNF4α directly activates multiple genes associated with the Classical subtype. HNF4α targets include genes that play a direct role in aspects of epithelial cell biology, such as *Lgals4*, which encodes a member of the Galectin family that mediates intercellular adhesion and apical trafficking (32), tight junction components like claudin proteins, and desmosomal proteins such as DSC2 and DSG2. Moreover, HNF4α activates other transcription factors (such as *Hnf1a*) that likely further amplify the transcriptional network governed by HNF4α. Second, HNF4α directly represses the expression of *Six1* and *Six4*, genes encoding homeodomain transcription factors that are highly expressed in the Basal-like subtype of PDAC and normally drive mesodermal development. Importantly, we find that HNF4α-negative PDAC cells are dependent on SIX4, and its close paralogue SIX1, for *in vitro* proliferation and *in vivo* tumorigenesis, suggesting that SIX4/1 activity may be a novel subtype-specific vulnerability in HNF4α-negative PDAC.

HNF4α is likely one of several endodermal lineage specifiers that regulate the Classical molecular subtype of PDAC (reviewed in (33)). For example, GATA6, FoxA1 and FoxA2 have been reported to promote epithelial differentiation and block EMT in human PDAC cell lines (34, 35). Moreover, GATA6 binds and activates *FOXA1* and *FOXA2* (34), and FoxA1/2 activate *HNF4A* and other endodermal lineage specifiers in multiple contexts (18, 36). Based on these data, a potential hierarchy of endodermal lineage specifiers that regulate the Classical subtype of PDAC begins to emerge. However, these studies did not formally compare gene expression changes induced by manipulation of each transcription factor with the Classical and Basal-like transcriptome profiles. It will likely be necessary to modulate the expression of these transcriptional regulators within the same physiologically relevant model system(s) to gain a comprehensive understanding of their individual impact on specification of the Classical subtype as well as their hierarchical regulatory relationships, which may involve extensive cross-talk and feedback loops as in normal tissue (37).

The ability of HNF4α to restrain tumor growth and overall progression of pancreatic tumors in our model systems is consistent with the better overall survival seen in the Classical subtype. Other endodermal cell fate determinants have been reported to restrain pancreatic cancer progression. For example, *Ptf1a* inhibits pancreatic tumorigenesis by not only preventing but also reversing tumor initiation (38). Nevertheless, it is clear the effect of endodermal lineage specifiers on tumor growth are to some degree context dependent. For example, an *in vitro* CRISPR screen identified HNF4α (as well as its target HNF1α) as dependencies in two out of nine human PDAC cell lines tested (39). In this regard, dichotomous effects on PDAC growth have been reported for HNF1α (40–42), PDX1 (43), FoxA1/2 (44–47) and GATA6 (34,48,49). It seems likely that even though the most malignant subset of PDAC downregulates the endodermal differentiation program, loss of this program may only confer a selective advantage in specific situations or stages of tumor progression. Furthermore, a dichotomous role for lineage specifiers in regulating malignant potential appears to be an emerging principle in cancer biology. For example, the pulmonary lineage specifier NKX2-1 can function as an oncogene in *EGFR*-mutant lung adenocarcinoma but restrains the growth of *KRAS*-mutant lung adenocarcinoma (50). Moreover, HNF4α itself is required for initiation and growth of NKX2-1-deficient lung adenocarcinoma, a subtype of lung cancer with a GI-like differentiation program similar to the Classical subtype of PDAC (18).

The transcriptional network governing the Basal-like subtype of PDAC is even less well understood than the Classical subtype. The transcription factors *Δ*Np63 (51, 52) and Gli2 (53) have been reported to promote growth and activate a Basal-like gene expression program in human cell lines. Loss of the histone demethylase KDM6A can lead to de-repression of *Δ*Np63 and drive the development of squamous and poorly differentiated PDAC (54). Our work identifies the homeodomain transcription factors SIX4 and SIX1 as HNF4α-repressed drivers of proliferation and growth in the Basal-like subtype. HNF4α binds directly to the promoters of *Six4* and *Six1* (as well as a potential distal regulatory element located between them), which is accompanied by decreased chromatin accessibility at these loci. The observed HNF4α chromatin binding and genomic accessibility changes correlated with a marked decreased proliferative capacity of PDAC cells. SIX1 and SIX4 are partially redundant drivers of mesodermal differentiation in tissues such as skeletal muscle, sensory neurons, kidneys and gonads (29). Several SIX family members have been implicated as having oncogenic functions in various types of cancer (55, 56), including SIX1 in PDAC (30, 31). However, SIX4 has not been previously shown to play a functional role PDAC, despite its elevated levels in the Basal-like subtype. Intriguingly, both SIX4 and SIX1 physically interact with co-factors that are potentially druggable targets. SIX4 physically interacts with the lysine demethylase UTX/KDM6A, which can be targeted by small molecule inhibitors (57). Moreover, the EYA family of transcriptional co-activators are essential for SIX1 function and harbor a druggable Tyr phosphatase activity (58, 59). Thus, SIX4 and SIX1 may represent targetable vulnerabilities of the basal subtype of PDAC. Additional studies are necessary to understand the mechanistic role of these transcription factors in the malignant progression of PDAC.

Taken together, our findings demonstrate the role of HNF4α as a regulator of a transcriptional network that restrains pancreatic cancer progression and imposes an epithelial cell identity and a Classical PDAC molecular subtype program that correlates with better outcome. Moreover, our results showed that HNF4α directly represses SIX1 and SIX4, which play an oncogenic function in HNF4α-negative PDAC, leading to the identification of potentially relevant therapeutic vulnerabilities in this highly aggressive form of the disease.

## Material and Methods

### Mice

Mice harboring *Kras^LSL-G12D^* (Jackson et al., 2001), *Kras^FSF-G12D^* (Young et al., 2011), *p53^flox^* (60), *Rosa26^LSL-tdTomato^* (18), *Hnf4a^flox^* (14), *Rosa-FSF-Cre^ERT2^* (61), *p53^frt^* (62), and *Pdx1-Cre* (19) alleles have been previously described. All animals were maintained on a mixed C57BL/6J x 129SvJ background. Animal studies were approved by the IACUC of the University of Utah and the University of California at San Francisco, and conducted in compliance with the Animal Welfare Act Regulations and other federal statutes relating to animals and experiments involving animals and adheres to the principles set forth in the Guide for the Care and Use of Laboratory Animals, National Research Council (PHS assurance registration number A-3031–01).

### Tamoxifen and Doxycycline Administration

Mice were fed *ad libitum* food pellets supplemented with tamoxifen (500 mg/ kg; Envigo, Indianopolis, IN) or doxycycline hyclate (625 mg/kg; Envigo, Indianopolis, IN) in place of standard chow starting 1 week before performing engraftments and for the entire duration of the experiments.

### Flank Tumor Transplantations

For subcutaneous allo- and xenografts experiments, were injected with 2.5 x10^5^ murine PDAC cells, 3.5 x10^5^ human PDAC cells or 5 x10^5^ PDAC cells from organoid cultures were resuspended in 50 μL of culture media, then mixed 1:1 with Matrigel (Corning). Cells were injected subcutaneously into the left or right flank NSG recipient mice. Subcutaneous tumor dimensions were measured with calipers twice weekly for the duration of the experiment, n = 8-10 independent tumors per group. Tumor volume was calculated using the standard modified ellipsoid formula ½ (Length x Width2) formula. After completion of the experiment, tumors were removed, weighed, and fixed for histological analysis.

### *In Vivo* Bioluminescence Imaging

Bioluminescence imaging was performed using an IVIS Spectrum *In Vivo* Imaging System (PerkinElmer). Mice bearing tumors of cell expressing firefly luciferase were injected with 200 μl/mouse of D-Luciferin (GoldBio) at 16.7 mg/ml. Optimal signals were obtained 10 min after subcutaneous injections of the D-Luciferin. Scanning was done with mice placed in prone position.

### Histology and Immunohistochemistry

All tissues were fixed in 10% formalin overnight and when necessary, lungs were perfused with formalin via the trachea. Organoids were first fixed in 10% formalin overnight and then mounted in HistoGel (Thermo Scientific, HG-4000-012). Mounted organoids and tissues were transferred to 70% ethanol, embedded in paraffin, and four-micrometer sections were cut. Immunohistochemistry (IHC) was performed manually on Sequenza slide staining racks (ThermoFisher Scientific, Waltham, MA). Sections were treated with Bloxall (Vector labs) followed by Horse serum 536 (Vector Labs, Burlingame, CA) or Rodent Block M (Biocare Medical, Pacheco, CA), primary antibody, and HRP-polymer-conjugated secondary antibody (anti-Rabbit, Goat and Rat from Vector Labs; anti-Mouse from Biocare. The slides were developed with Impact DAB (Vector) and counterstained with hematoxylin. Slides were stained with antibodies to BrdU (BU1/75, Abcam, Cambridge, MA) 1:400, HA tag (C29F4, CST) 1:400, Galectin 4 (AF2128, R&D Systems) 1:200, HNF4α (C11F12, CST) 1:500, PK-LR (EPR11093P, Abcam) 1:500, SIX1 (D5S2S, CST) 1:100, HNF4α P1 and P2 (Human HNF-4 alpha/NR2A1 MAb (Clone K9218) PP-K9218-00, Human HNF-4 alpha/NR2A1 MAb (Clone H6939) PP-H6939-00) 1:100. Pictures were taken on a Nikon Eclipse Ni-U microscope with a DS-Ri2 camera and NIS-Elements software. Mitoses quantitation and histological analyses were performed on hematoxylin and eosin-stained or IHC-stained slides using NIS-Elements software. All histopathologic analysis was performed by a board-certified anatomic pathologist (E.L.S.).

### Derivation of 2D Cell Lines and 3D Organoid Cultures

Cell lines and 3D cultures were created by enzymatic and mechanical dissociation of pancreatic tumors or normal pancreatic tissue harvested from *Kras^LSL-G12D^; p53^flox/+^; Hnf4a^flox/flox^*; Pdx1-Cre and *Kras^FSF-G12D^; p53^fr/frt^; Hnf4a^F/F^; RosaCre^ERT2^* mice, respectively. Two dimensional cultures were grown in DMEM supplemented with 10%FBS. For organoid derivation, single cell suspensions were transduced with an adenovirus expressing Flp recombinase for 2 hs at 37°C, washed, and seeded in grow factor-reduced Matrigel (Corning) and grown in 50% L-WRN conditioned media (63). HNF4α expression restoration was induced by supplementing the corresponding media with 1ug/ml doxycycline (Sigma-Aldrich, D9891) and *Hnf4a* genetic loss was carried out by adding 4-OH-Tamoxifen (Cayman Chemical, 68392-35-8) to the culture media.

### Cell proliferation assay

We used a live cell imaging system (IncuCyte) to directly measure cell proliferation while monitoring morphological changes over time. Number of cells was tracked by detecting the tdTomato present in HC569 and HC800 cell lines or by transducing cells to express eGFP fused to the histone H2B, which labels cell nuclei.

### Vector Cloning

pCW-TRE-HNF4α lentiviral vector was generated by PCR amplifying the HNF4α isoform 8 cDNA from murine mucinous adenocarcinomas tumors and cloning into HpaI-PacI sites of pCW22-TRE-cDNA;Ubc-rtTA plasmid. Correct identity and orientation of the construct was confirmed via Sanger sequencing.

### Lentiviral Production

Lentivirus and retrovirus were produced by transfection of 293T cells with TransIT-293 (Mirus Bio). Packaging vectors Δ8.9 (gag/pol) and CMV-VSV-G (DuPage et al., 2009) were used for lentiviral production. Supernatant was collected at 48, 60 and 72 hs post-transfection, centrifuged, and filtered using 0.45um filter units.

### cDNA Synthesis and qRT-PCR

Quantitative RT–PCR was performed on Trizol-extracted RNA using the iScript Reverse Transcription Supermix (BIO-RAD). qPCR reactions were performed using SsoAdvanced Universal Probes Supermix (BIO-RAD) and a CFX384 Touch Real-Time PCR Detection System (BIO-RAD). The following Taqman probes were used: Hnf4a (Mm01247712_m1) and Pklr (Mm00443090_m1). Transcript levels were normalized to Ppia (Mm02342430_g1) and quantitated by the ΔΔCt method.

### Immunoblotting

Cells were lysed on ice for 20 minutes in RIPA buffer (50mM Tris HCl pH 7.4, 150 mM NaCl, 0.1 % (w/v) SDS, 0.5% (w/v) sodium deoxycholate, 1% (v/v) Triton X-100) plus Complete protease phosphatase inhibitor cocktail (A32961, ThermosFiher Scientific). Cells were pelleted for 10 minutes at 4 °C and protein concentration was quantitated with the Coomassie (Bradford) Protein Assay (ThermoFisher Scientific). Lysates were separated on Tris-Glycine (TGX) precast gels (Biorad) and transferred to nitrocellulose membranes (ThermoFisher Scientific). Membranes were probed with antibodies to HA tag (C29F4, CST, 1:5000), HNF4α (C11F12, CST, 1:2000), ß-Tubulin (DSHB, 1:2000), Vinculin (Abcam, 1:20000), Six1 (D5S2S, CST, 1:1000), Six4 (D-5, Santa Cruz Biotechnology, 1:1000), and the appropriate species conjugated secondary antibodies (ThermoFisher Scientific, 1:20000), and visualized using an Odyssey CLx Imaging System (LI-COR Biosciences).

### Chromatin Immunoprecipitation (ChIP-seq) Assay and Analysis

After treating HC569 and HC800 with DOX or vehicle for 1 week, cells were crosslinked with 1% formaldehyde for 10 min at room temperature and then treated with 125 mM glycine for 5 min to stop crosslinking. Crosslinked cells were then washed with cold PBS and scraped to harvest. ChIP was performed as previously described (Reddy et al., 2009). Sonication was performed on Active Motif EpiShear Probe Sonicator with 8 cycles of 30 s of amplitude 40% sonication and 30 s of rest. The antibody used was the purified anti-HA.11 Epitope Tag Antibody (16B12, BioLegend). Reads were aligned to the mm10 build of the mouse genome using Bowtie with the following parameters: -m 1 -t–best -q -S -l 32 -e 80 -n 2 (64). Peaks were called using Model-Based Analysis of ChIP-seq-2 (MACS2) (Zhang et al., 2008) with a p value cutoff of 1e—10 and the mfold parameter constrained between 15 and 100. ChIP-seq peaks that were replicated within each cell line were used for downstream analysis. Motif finding was performed on 100 bp surrounding the top 1000 peaks based on their integer score (column 5 of narrowPeak file). Motifs were discovered using the meme suite (Bailey et al., 2009), searching for motifs between 6 and 50 bases in length, with zero or one occurrence per sequence. Genes were assigned to ChIP-seq peaks using GREAT with a 100kb distance cutoff (65).

### Assay for Transposase Accessible Chromatin (ATAC-seq) and Analysis

Following doxycycline treatment for 1 weeks, ATAC-seq was performed on 250,000 cells for each library (described by Buenrostro et al., 2013). Tn5 transposase, with Illumina adapters, was constructed as outlined earlier (Picelli et al., 2014). Sequencing reads were aligned to mm10 using Bowtie (Langmead et al., 2009) with the following parameters: -m 1 -t–best -q -S -l 32 -e 80 -n 2. SAM files were converted to BAM files and sorted using samtools (Li et al., 2009). MACS2 (Zhang et al., 2008) was used to call peaks without a control input. We used a cutoff p-value of 1e-10 when calling peaks. Feature Counts (Liao et al., 2014) was used to quantify reads that aligned in regions ± 250 bp Hnf4α-binding site summits from the ChIP-seq experiments. These reads were then normalized and differential expression was determined by comparing samples in a pairwise manner using the DESeq2 package for R (Love et al., 2014). DESeq2 was used with a multivariate model that corrected for cell line and then identified significant effects from doxycycline treatment. Genes were assigned to ATAC-seq peaks using GREAT with a 100kb distance cutoff (65).

### RNA sequencing

RNA was isolated from aniline blue-stained FFPE sections of micro-dissected autochthonous tumors by the Molecular Diagnostic core facility (HCI) using the miRNAeasy kit (Qiagen). IHC for HNF4α on serial sections was used to identify viable tumor areas positive or negative for HNF4α. RNA was isolated from cell lines and organoid cultures by Trizol-chloroform extraction followed by column-based purification. The aqueous phase was brought to a final concentration of 50% ethanol, and RNA was purified using the PureLink RNA Mini kit according to the manufacturer’s instructions (ThermoFisher Scientific). Library preparation was performed using the TruSeq Stranded RNA kit with Ribo-Zero Gold (Illumina). Libraries from FFPE purified samples were sequenced on an Illumina HiSeq 2500 (50 cycle single-read sequencing) and samples from 2D cells and organoids were sequenced on a NovaSeq 6000 (50bp paired-end sequencing).

### RNA-seq Data Processing and Analysis

Mouse FASTA and GTF files were downloaded from Ensembl release 92 and a reference database was created using using STAR version 2.5.4a (66)) with splice junctions optimized for 50 base pair reads. Reads were trimmed of adapters using cutadapt 1.16 (67) and then aligned to the reference database using STAR in two pass mode to output a BAM file sorted by coordinates. Mapped reads were assigned to annotated genes in the GTF file using featureCounts version 1.6.3 (68). The gene counts were filtered to remove 18,611 features with zero counts and 12,997 features with fewer than 5 reads in every sample. Differentially expressed genes were identified using a 5% false discovery rate with DESeq2 version 1.22.2 (69).

### Comparison of RNA-seq Data with Public Data

TCGA survival data was downloaded from Supplemental Table 1 of (5) while the mRNA expression data was downloaded using the R package *TCGABiolinks*. The cohort was trimmed to only include samples of high purity confirmed adenocarcinomas (>30% tumor, n = 76), and then was further subdivided by HNF4A mRNA expression. The PACA-AU data and associated metadata were downloaded from the icgc.org data portal, with the supplemental data coming from (2). This study excluded samples with <40% purity, so no further trimming was necessary. For generation of basal-like and classical scores of mouse tumors, the human gene signatures from Moffitt et al. were used (n = 25 per subtype). Only genes with identified mouse homologs were considered (basal-like n = 14, classical n = 18). Expression was log normalized, and the mean expression of all gene set members was defined as the overall expression score. For gene set enrichment, expression data was split into two conditions (Hnf4a+ and Hnf4a-) and a rank order list prepared per gene using the difference in means divided by the sum of standard deviations for samples from each condition.

For presentation of genes that are differentially expressed across model systems (Supplemental Table S5), “up” genes includes all three datasets, whereas “down” genes includes only autochthonous tumors and 2D cell lines because there was little overlap between “down” genes in the organoid system in comparison with the other models.

### CRISPR-Cas9 Genome Editing

SIX1 and SIX4 knock-out cell lines were generated by CRISPR-Cas9. LentiCRISPRv2 vectors, which express the single guide RNA (sgRNA), Cas9 and a selection cassette, were obtained from Addgene (ID: 52961 and 98291).

Cells were transduced and then subjected to puromycin or hygromycin selection, as corresponds. Two different sgRNA sequences were used to target *Six1*, four were tested for *Six4*, and an inert non-targeting guide was used as a control.

sgSIX1 GTGGCTGAAAGCGCACTACG (Reference: (31)). sgSIX1#2 GGAGGGAACCTGGAACGCCT sgSIX4#1 AAGTGCGGCGGACATCAAGC sgSIX4#2 CGCGGCGTCCCCCGGCTCCA sgSIX4#3 GCGCACCGAGAAGTGGCGGG sgSIX4#4 TGCCTCCAGGGTAAGCCGGG sgNT GCGAGGTATTCGGCTCCGCG (Reference: (70)).

Individual cells were allowed to expand for a period of 3–6 weeks, after which time clonal expansions of Six1 knock outs grew. Confirmation of efficient deletion was carried out by detecting protein loss by Immunoblot for endogenous SIX1 and exogenously over-expressed SIX4, as its endogenous levels were essentially undetectable due to lack of antibody sensitivity.

### Statistics

Statistical analyses were carried out using Prism 8. p-Values were calculated using Mann-Whitney U test, Chi-square test, Log-rank test or Wilcoxon test. RNA-Seq statistics are described above.

## Supporting information

Supplemental figures with legends

## Data availability

Gene expression, ChIP-Seq and ATAC-seq data are accessible in the NCBI Gene Expression Omnibus (GEO) database under accession number GSE138145 and GSE138463.

## Acknowledgements

We are grateful to members of the Snyder lab for suggestions and comments. We thank Brian Dalley for sequencing expertise and James Marvin for FACS expertise. Core facilities (PRR, BMP, DNA sequencing, Genomics/Bioinformatics, Flow Cytometry). Research reported in this publication utilized shared resources (including Flow Cytometry, High Throughput Genomics, Bioinformatics, and Biorepository and Molecular Pathology) at the University of Utah and was supported by the National Cancer Institute of the National Institutes of Health under Award Number 5R21CA194764-03. Work in the flow cytometry core was also supported by the National Center for Research Resources of the National Institutes of Health under Award Number 1S20RR026802-1. ELS was supported in part by a Career Award for Medical Scientists from the Burroughs Wellcome Fund, a V Scholar Award, the NIH (R01CA212415) and institutional funds (Department of Pathology and Huntsman Cancer Institute, University of Utah).

## Author contributions

SC and ES designed experiments. SC, VB, KB and HC performed experiments. SC, LTH, JV, CS, RM, JG and ES analyzed data. ES performed histopathologic review. SC, and ES wrote the manuscript. All authors discussed results, reviewed and revised the manuscript.

