## Supplemental figures with legends for "Reciprocal regulation of pancreatic ductal adenocarcinoma growth and molecular subtype by HNF4α and SIX1/4"

**Camolotto et al.**

**Supplemental Figures**

Figure S1, Camolotto et al.

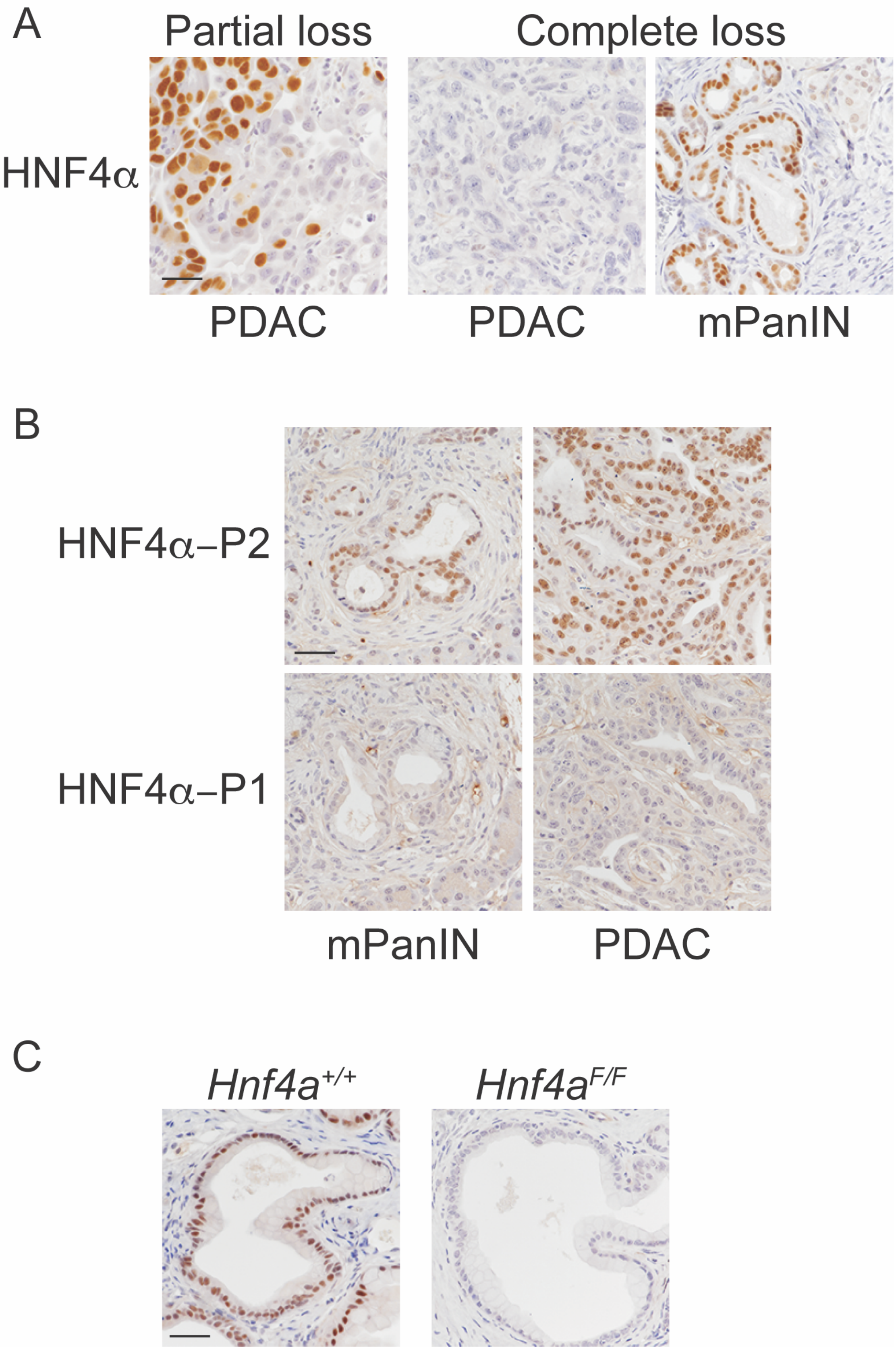

**Figure S1, related to figure 1**.

A. IHC demonstrating partial (left) and complete (center) stochastic HNF4α loss in PDAC arising in *Kras*$^{LSL\text{-}G12D/+;}$ *p53*$^{F/+}$*; Pdx1-Cre; Hnf4a*$^{+/+}$ mice (12 weeks of age). Right: HNF4α-positive PanIN from pancreas with HNF4α-negative PDAC. Scale bar: 100 microns.

B. IHC for P1 and P2 isoforms of HNF4α in pancreatic neoplasia from *Kras*$^{LSL\text{-}G12D/+;}$ *p53*$^{F/+}$*; Pdx1-Cre; Hnf4a*$^{+/+}$ mice (12 weeks of age). mPanIN: Pancreatic intraepithelial neoplasia. Scale bar: 100 microns.

C. IHC for HNF4α on pancreatic neoplasia from *Kras*$^{LSL\text{-}G12D/+;}$ *p53*$^{+/+}$*; Pdx1-Cre; Hnf4a*$^{F/F}$ mice and *Hnf4a*$^{+/+}$ controls at 9 months of age. Scale bar: 100 microns.

Figure S2, Camolotto et al.

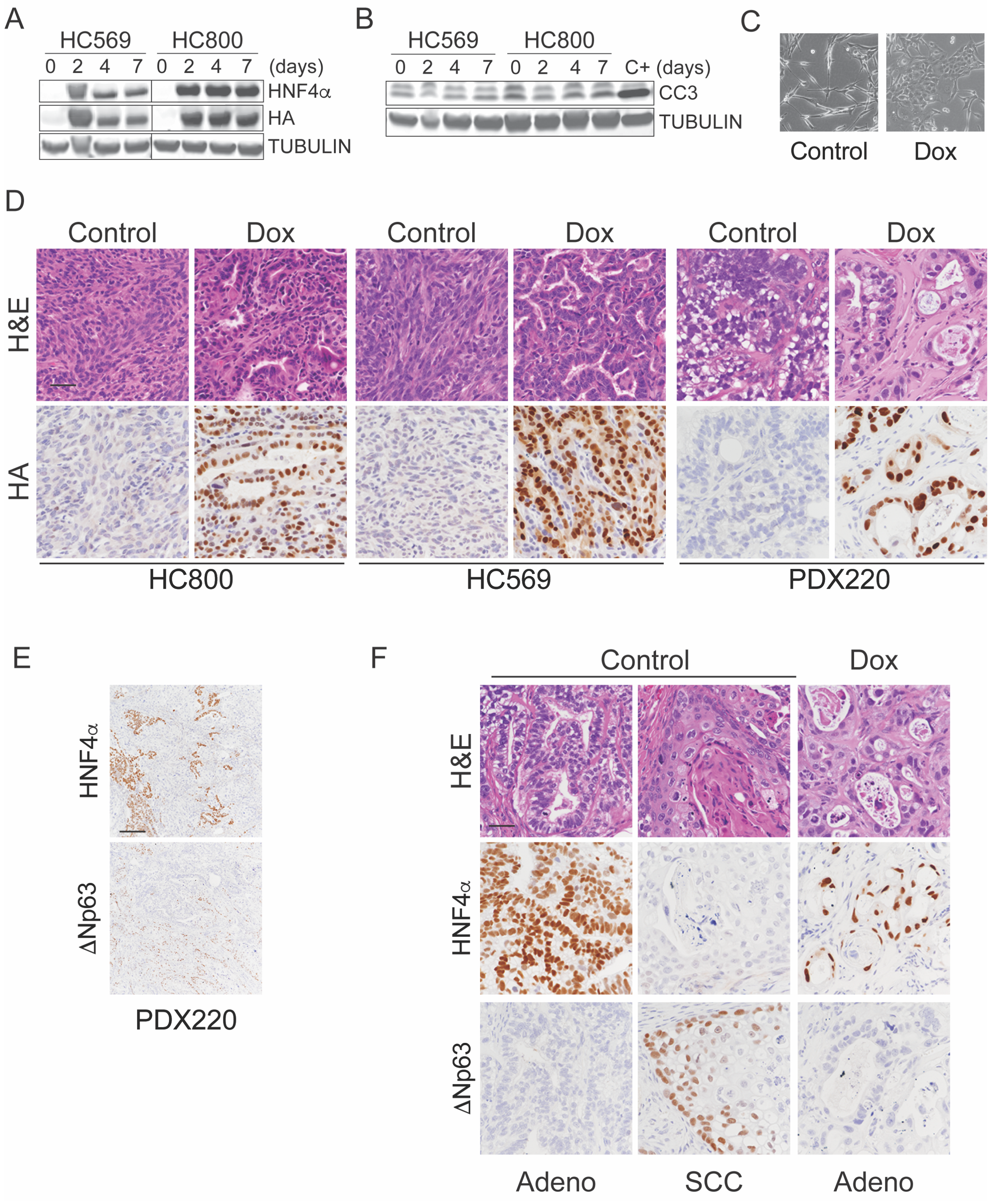

**Figure S2, related to figure 2**.

A-B. Immunoblot analysis for indicated proteins on two independent murine cell lines (HC800 and HC569) from PDAC arising in *Kras*$^{LSL\text{-}G12D/+;}$ *p53*$^{F/+}$*; Pdx1-Cre; Hnf4a*$^{F/F}$ mice. Cells were transduced with lentivirus encoding doxycycline-inducible HA-tagged HNF4$\alpha$8 isoform and then selected with blasticidin. Cells were treated with doxycycline for 2, 4 or 7 days prior to lysis.

C. Bright field images of HC800 cell lines treated with doxycycline or vehicle for 72hs. HC800 cell morphology changes from a spindle to a classic epithelial cobblestone growth pattern with HNF4$\alpha$ restoration.

D. H&E and IHC for HA on subcutaneous tumors from HC800, HC569 and PDX220 cells in the presence or absence of doxycycline treatment. Scale bar: 100 microns.

E. IHC for HNF4$\alpha$ for PDX220 cells subcutaneously injected in NSG mice. Scale bar: 500 microns.

F. H&E and IHC for HNF4$\alpha$ and $\Delta$Np63 on subcutaneous tumors from PDX220 cells in the presence or absence of doxycycline treatment. Scale bar: 100 microns.

Figure S3, Camolotto et al.

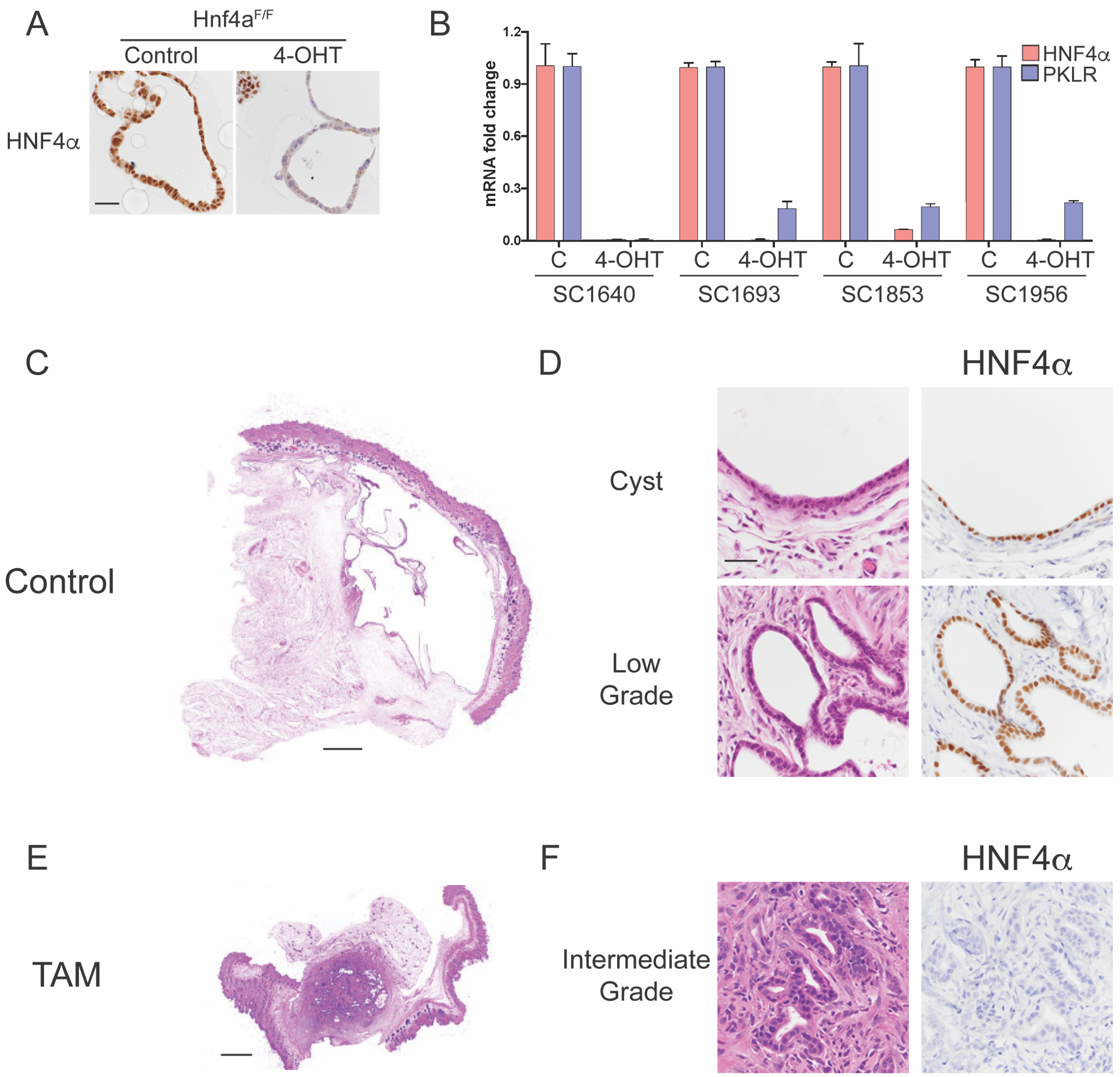

**Figure S3, related to figure 3**

A. Representative IHC for HNF4$\alpha$ in PDAC organoid cultures derived from pancreata of *Kras*$^{FSF-G12D/+}$; *p53*$^{Frt/Frt}$; *Rosa26*$^{FSF-CreERT2}$; *Hnf4a*$^{F/F}$ mice. Cells were treated with ethanol (control) or 4-hydroxytamoxifen (4-OHT) to activate Cre$^{ERT2}$ and delete *Hnf4a*. Scale bar: 100 microns.

B. qRT-PCR in PDAC organoids cultures for indicated transcripts. Cells were treated with ethanol (Control) or 4-hydroxytamoxifen (4-OHT) to activate Cre$^{ERT2}$ and delete *Hnf4a*.

C-F. H&E and IHC analysis of SC1693 organoids injected subcutaneously into NSG mice. Mice were fed control (C-D) or tamoxifen (E-F) chow starting 1 week prior to flank injection. Tumors were analyzed at 6 weeks post injection. Left images: scanning magnification, scale bar: 1 mm. Right: High power images, scale bar: 100 microns.

Camolotto et al., Figure S4

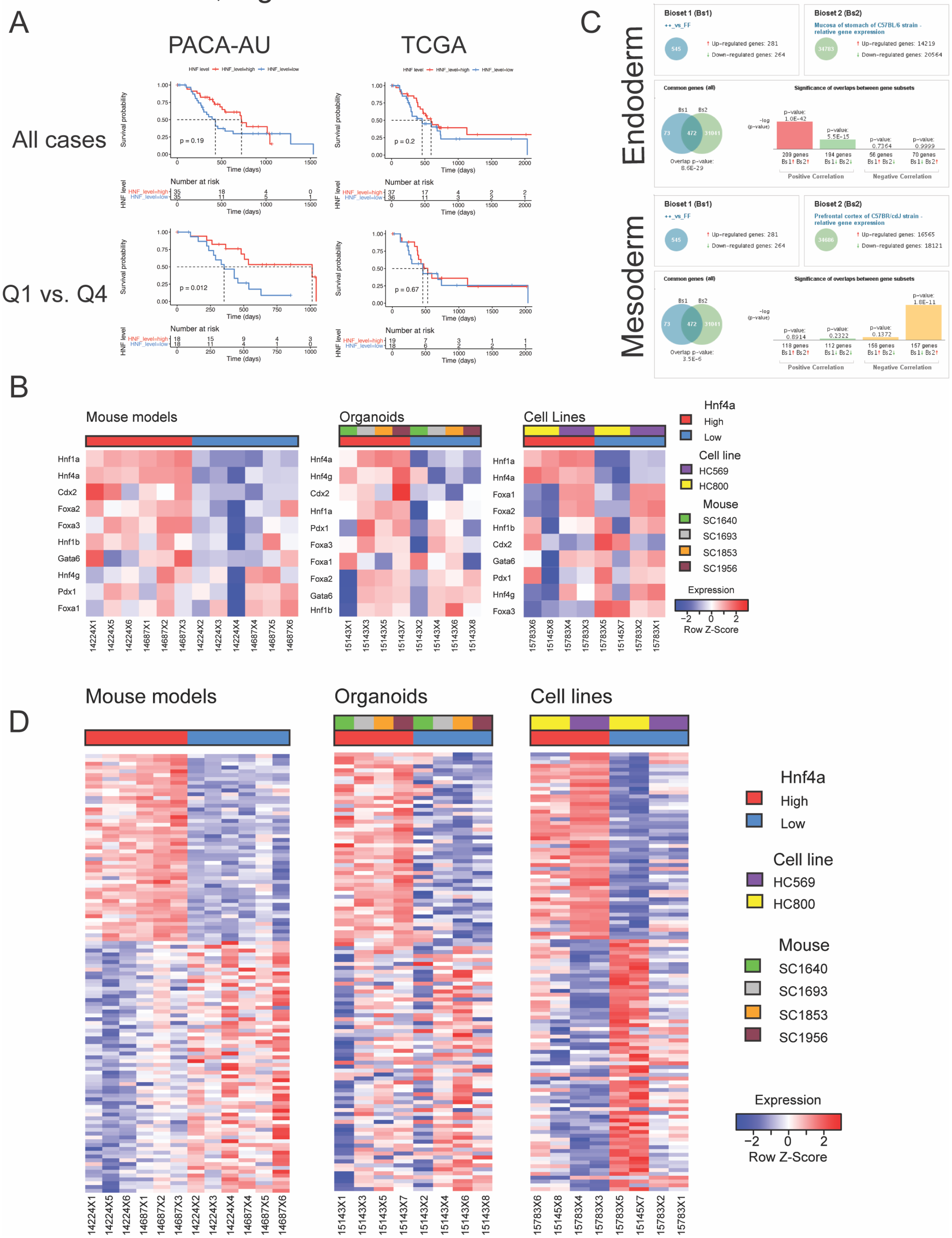

**Figure S4, related to figure 4**.

A. Survival analysis of patients with HNF4A high vs low PDAC in PACA-AU (left) and TCGA (right) cohorts. Top row: comparison of upper half vs. lower half (split around median HNF4A expression). Bottom row: comparison of upper vs. lower quartile. Analysis was limited to high purity tumors (>30% cellularity).

B. Heatmaps showing relative expression of endodermal lineage markers as measured by RNA-Seq in PDAC models. Samples sorted by HNF4α expression, rows clustered by Pearson.

C. Representative tissues exhibiting positive and negative correlations with HNF4α expression in autochthonous tumors (Body Atlas function, Illumina Correlation Engine).

D. Heatmaps showing relative expression of genes identified by DESeq2 to be consistently altered along with *Hnf4a* modification. Samples sorted by t statistic.

Camolotto et al., Figure S5

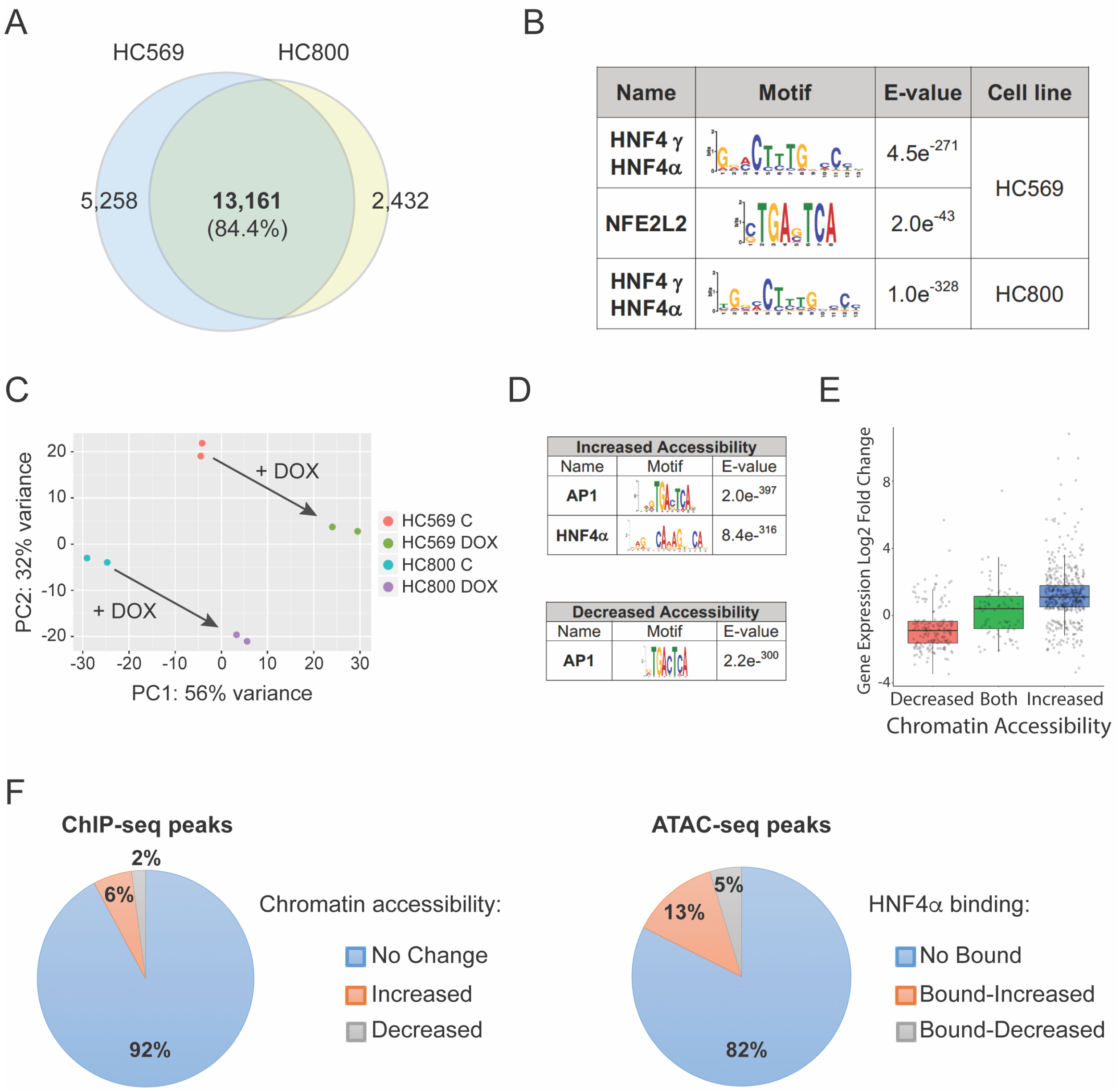

**Figure S5, related to figure 5**.

A. Venn diagram shows the overlap between HNF4α ChIP-seq peaks identified in HC569 and HC800 cells.

B. Motif enrichment in HNF4α binding sites. STAMP logos and enrichment statistics are shown for both HC569 and HC800 cells.

C. Principal component analysis of ATAC-seq peaks within control and dox-treated HC569 and HC800 cells. Each dot represents an individual sample.

D. Significantly enriched transcription factor motifs in regions with increased and decreased chromatin accessibility.

E. Boxplot shows distribution of gene expression level among ATAC-seq regions with different chromatin accessibility.

F. Pie charts displaying distribution of HNF4α bound genomic regions with differential chromatin accessibility (left panel) and distribution of differentially accessible chromatin regions bound or not to HNF4α (right panel).

### Camolotto et al., Figure S6

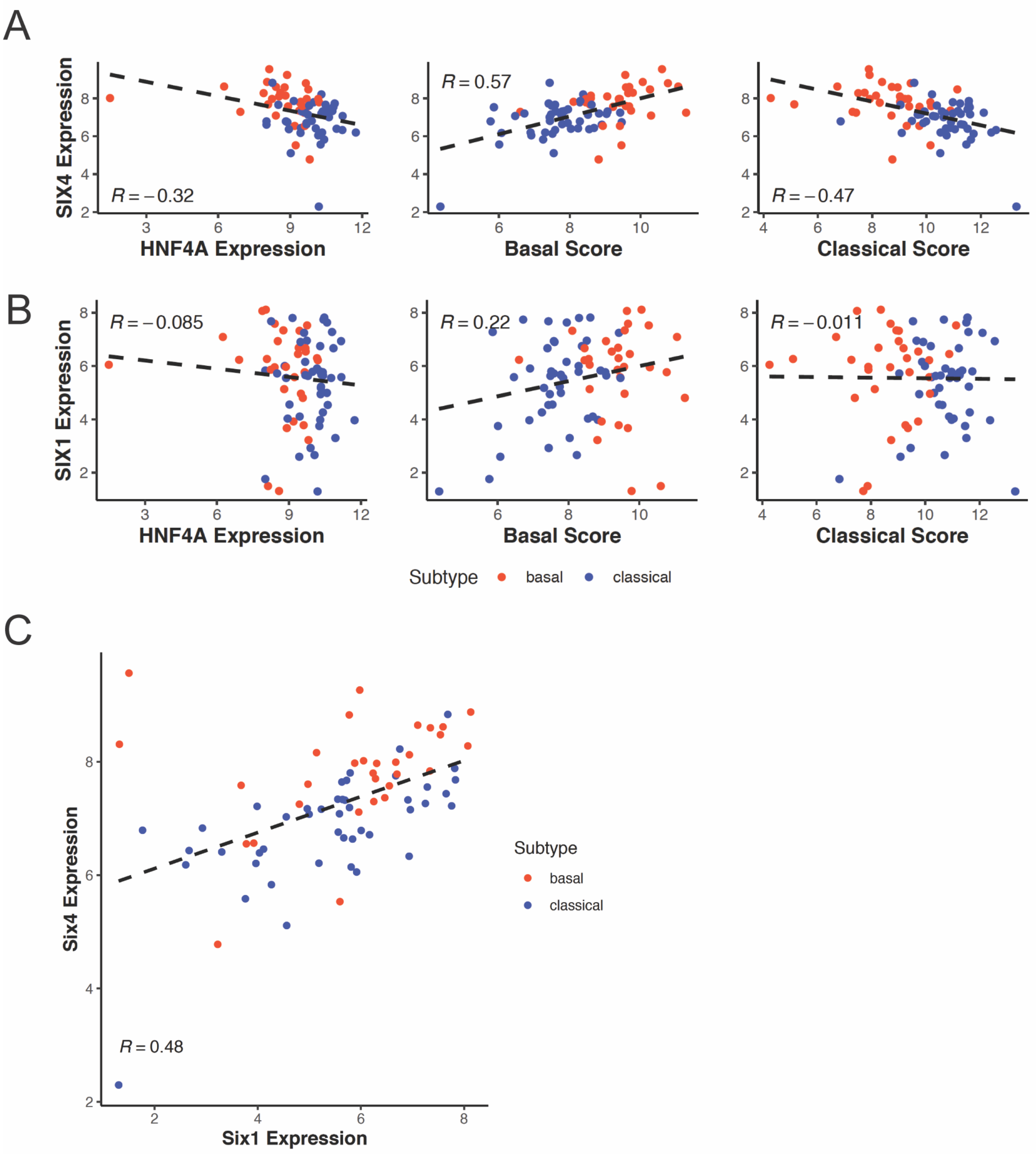

**Figure S6, related to Figure 6**

A. Correlation between SIX4 and HNF4A, Classical score and Basal-like score in human PDAC (high purity cases, TCGA).

B. Correlation between SIX1 and HNF4A, Classical score and Basal-like score in human PDAC (high purity cases, TCGA).

C. Correlation between SIX4 and SIX1 in human PDAC (high purity PDAC, TCGA).

Camolotto et al., Figure S7

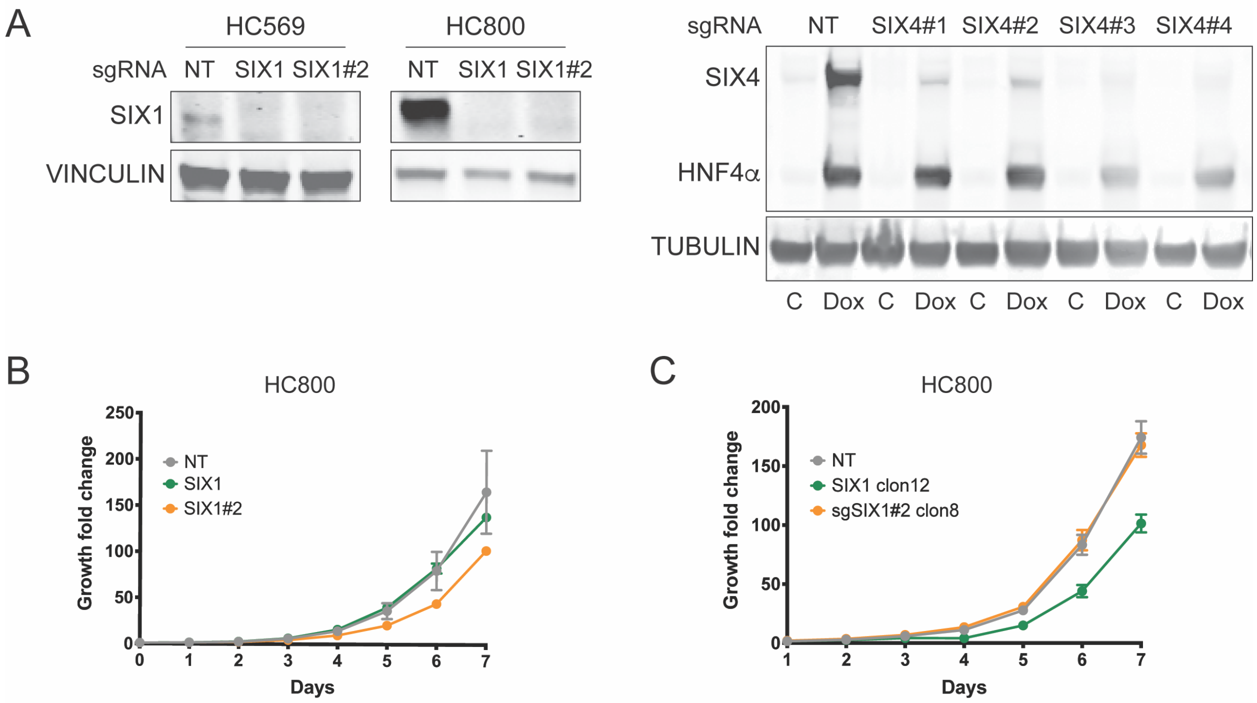

**Figure S7, related to Figure 7**

A. CRISPR/Cas9 validation. Immunoblot for SIX1 in HC569 and HC800 expressing Cas9 and either of the 2 indicated sgRNA sequences targeting *Six1* (left panel). Immunoblot for SIX4 in HC800 overexpressing doxycycline inducible SIX4 and HNF4$\alpha$. The 4 indicated sgRNA sequences recognizing *Six4* were tested (left panel).

B. Quantitation of proliferation in HC800 cells stably expressing Cas9 and the indicated sgRNA against *Six1* (n= 3 biological replicates). Representative growth curves of two independent experiments with similar results are shown. Data represents mean +/- SEM. $p \leq 0.05$, Wilcoxon test.

C. Quantitation of proliferation in single cell cloned HC800 cells stably expressing Cas9 and the indicated sgRNA against *Six1* (n= 3 biological replicates, 2 independent experiments). Graph represents mean +/- SEM. $p \leq 0.05$, Wilcoxon test.
